# Lipidomic Profiling Reveals an Age-Related Deficiency of Skeletal Muscle Proresolving Mediators that Contributes to Maladaptive Tissue Remodeling

**DOI:** 10.1101/2020.11.05.370056

**Authors:** James F. Markworth, Lemuel A. Brown, Eunice Lim, Jesus A. Castor-Macias, Jacqueline Larouche, Peter C. D. Macpherson, Carol Davis, Carlos A. Aguilar, Krishna Rao Maddipati, Susan V. Brooks

**Author notes:** Susan V. Brooks, PhD, Christin Carter-Su Collegiate Professor of Physiology, Professor of Biomedical Engineering, Professor of Molecular & Integrative Physiology, University of Michigan, 2029 BSRB, 109 Zina Pitcher Pl. Ann Arbor, MI 48109-2002.

## Abstract

Chronic inflammation and deregulated acute immune cell responses to injury contribute to age-associated skeletal muscle dysfunction. Specialized pro-resolving mediators (SPMs) control inflammation and support myofiber regeneration in young mice, but their role in aging muscle remains unknown. Here we examined the effect of age on the mediator lipidome of skeletal muscle via LC-MS based lipidomic profiling and tested whether systemic administration of the SPM resolvin D1 (RvD1) could limit excessive inflammation and improve the regenerative capacity of aged muscle. Aged mice displayed chronic low-grade muscle inflammation prior to injury and this was associated with a basal deficiency of lipoxygenase (LOX) derived SPMs as well as anti-inflammatory cytochrome P450 (CYP) derived lipid epoxides. Following muscle damage, young and aged mice produced similar amounts of pro-inflammatory cyclooxygenase (COX) and 12-LOX metabolites, but aged mice mounted a markedly deficient SPM response. This was associated with heightened leukocyte recruitment, impaired myofiber regeneration, and delayed recovery of strength. Systemic treatment with RvD1 had minimal impact on excessive myeloid cell infiltration and defective myofiber regeneration in aged mice. Nevertheless, RvD1 treatment did suppress inflammatory cytokines, modulated muscle stem cells, limited maladaptive tissue remodeling, and improved recovery of specific muscle force. We conclude that aging results in a marked deficiency of local SPM biosynthesis within muscle and that immunoresolvents may be attractive novel therapeutics for the treatment of muscular injuries and associated pain in the elderly, due to positive effects on recovery of muscle function without the negative side effects on myofiber regeneration of traditional anti-inflammatory treatments.

## Introduction

Aging results in a progressive decline in skeletal muscle mass and function that contributes to frailty, loss of mobility, and increased mortality in the elderly (Faulkner, Larkin, Claflin, & Brooks, 2007). Aged muscles are also more susceptible to myofiber damage and have a reduced ability to regenerate successfully if injured (Blau, Cosgrove, & Ho, 2015). Potential sources of this dysfunction include an accumulation of macrophages (MΦ) within resting muscle tissue in advanced age (Wang, Wehling-Henricks, Samengo, & Tidball, 2015), as well as transient dysregulation of local acute myeloid cell responses to muscle injury (Sloboda, Brown, & Brooks, 2018). On this basis, targeting the immune system has shown promise to rejuvenate the regenerative capacity of aged muscle (Kuswanto et al., 2016; Oh et al., 2016; Patsalos et al., 2018) or even limit age-associated muscle wasting (Kim et al., 2020; Rieu et al., 2009; Wang et al., 2015; Wang et al., 2019; Wang, Welc, Wehling-Henricks, & Tidball, 2018).

The immune response to skeletal muscle injury begins with rapid infiltration of polymorphonuclear leukocytes (PMNs) (Orimo, Hiyamuta, Arahata, & Sugita, 1991), followed by an appearance of blood monocyte-derived MΦ (Tidball, Berchenko, & Frenette, 1999). These MΦ initially exhibit a pro-inflammatory phagocytic phenotype, but then later switch to an anti-inflammatory reparative subset that function to resolve inflammation and support ensuing muscle regeneration (Arnold et al., 2007). In general, the onset of the resolution phase of the acute inflammatory response is actively controlled by endogenous specialized pro-resolving lipid mediators (SPMs) including the lipoxins (Serhan, Hamberg, & Samuelsson, 1984), E-series (Serhan et al., 2000) and D-series resolvins (Serhan et al., 2002), protectins (Mukherjee, Marcheselli, Serhan, & Bazan, 2004), and maresins (Serhan et al. 2009). Each SPM family is enzymatically derived from essential dietary polyunsaturated fatty acid (PUFA) substrates via the coordinated actions of the 5-12-, and 15- lipoxygenase (LOX) pathways and share defining actions to inhibit further PMN recruitment to the site of inflammation, while stimulating MΦ functions necessary for timely resolution (Serhan & Levy, 2018). SPMs are produced in response to muscle injury in both rodents (Giannakis et al., 2019; Markworth et al., 2020) and humans (Gangemi et al., 2003; Markworth et al., 2013; Vella et al., 2019), suggesting that they may play an important role in the immunological and adaptive responses to myofiber damage (Markworth, Maddipati, & Cameron-Smith, 2016).

Current approaches to clinical management of muscle injuries focus predominantly on blocking production of early pro-inflammatory mediators. Non-steroidal anti-inflammatory drugs (NSAIDs) lessen the harmful effects of muscle injury in the short term by reducing strength loss and muscle soreness (Morelli, Brown, & Warren, 2017); however, NSAIDs also delay timely resolution of the acute inflammatory response and can impair myofiber growth and regeneration (Markworth et al., 2016). In contrast to NSAIDs, therapeutic administration of resolution agonists such as native SPMs or drug mimetics, termed immunoresolvents, can actually improve muscle regenerative outcomes in young mice and may thus offer an attractive alternative to classical anti-inflammatory approaches to the clinical management of muscle injuries (Giannakis et al., 2019; Markworth et al., 2020).

Recent studies have suggested that aging is associated with an overabundance of pro-inflammatory eicosanoids (e.g. prostaglandins) and/or a deficiency of SPMs (e.g. resolvins) (Arnardottir, Dalli, Colas, Shinohara, & Serhan, 2014; Gangemi et al., 2005; Halade, Kain, Black, Prabhu, & Ingle, 2016; Kain et al., 2019; Lopez et al., 2015; Rymut et al., 2020). Indeed, mice lacking one key SPM receptor (ALX/FPR2) develop a premature aging phenotype characterized by obesity, reduced life span, chronic low-grade inflammation, and dysregulated acute immune cell responses to an inflammatory challenge (Tourki et al., 2020). Treatment of aged mice with resolvins has been shown to limit excessive PMN influx and stimulate PMN clearance in a murine model of peritonitis (Arnardottir et al., 2014). Moreover, aged mice also displayed excessive PMN infiltration of the lung following remote ischemia-reperfusion injury and this response was blunted by intravenous administration of the native SPM resolvin D1 (RvD1) (Rymut et al., 2020). However, the endogenous role of SPMs and potential therapeutic applicability of immunoresolvents in the context of skeletal muscle aging has not yet been examined.

Therefore, in the current study we investigated the effect of aging on the mediator lipidome of resting skeletal muscle as well as dynamic shifts in local muscle lipid mediator profile in response to sterile skeletal muscle injury. Furthermore, we examined whether daily systemic treatment with RvD1 as an immunoresolvent could potentially limit inflammation and stimulate muscle tissue regenerative responses in aged mice.

## Results

### Age-Associated Loss of Muscle Mass and Strength

Aged mice (26-28 mo) had similar body weights, but lower tibialis anterior (TA) muscle mass than young mice (4-6 mo) (Figure 1A). Maximal *in-situ* nerve-stimulated TA isometric force production (P_o_) was lower in aged mice and this deficit persisted after accounting for their smaller muscle size (specific P_o_, sP_o_) (Figure 1B). TA muscles from aged mice contained similar numbers of total muscle fibers, but had a substantial reduction in mean myofiber cross-sectional area (CSA) (Figure 1C & D). Aging also resulted in a loss of type IIa and IIx myofibers from the TA, an increased presence of IIb fibers, and a reduction in mean CSA for all type II fibers (IIa, IIx and IIb) (Figure 1C & E). These data show that these mice developed a robust sarcopenic phenotype in advanced age.

**Figure 1.**
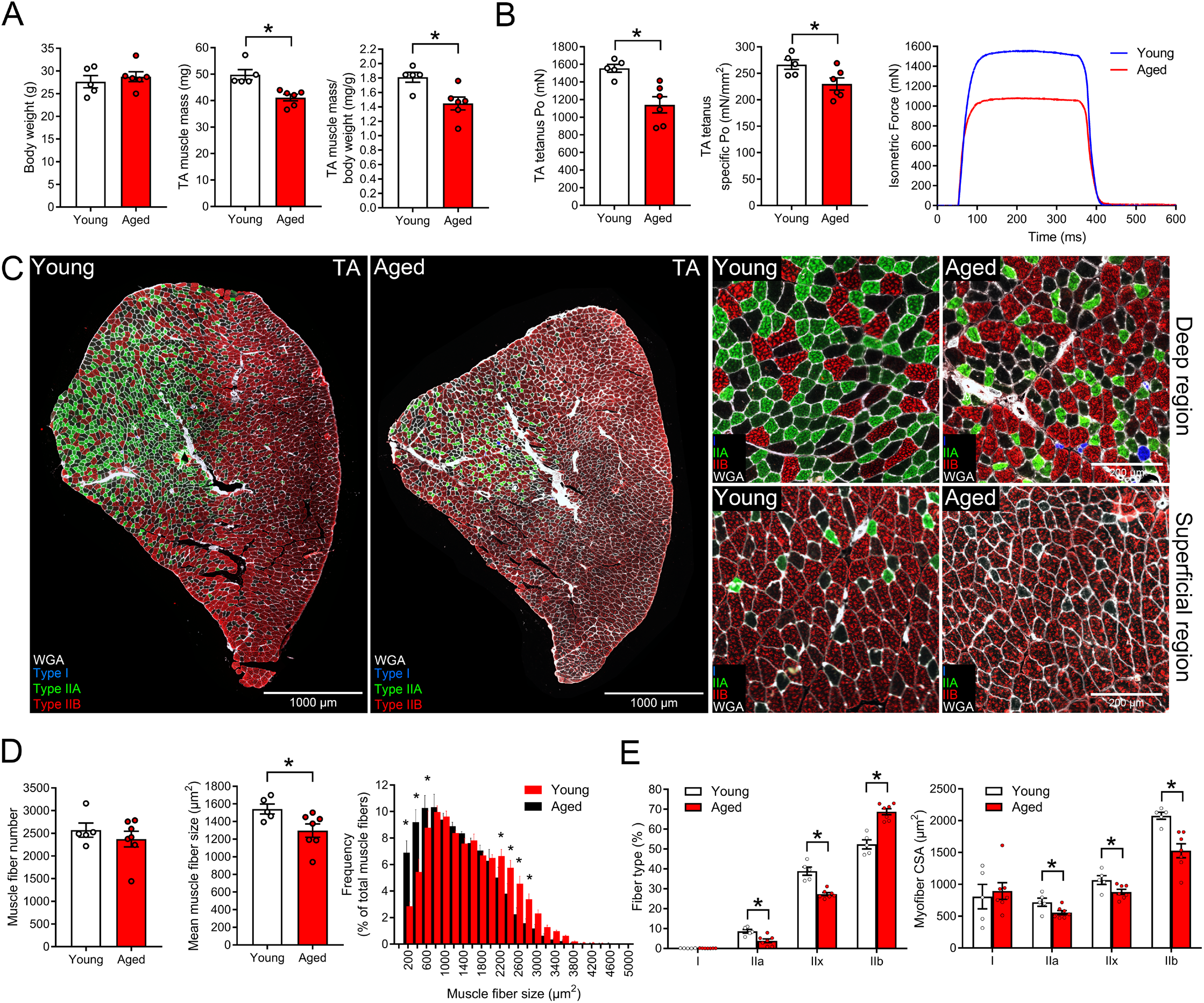
Age-associated basal skeletal muscle dysfunction: A: Body weight (g), absolute tibialis anterior (TA) muscle mass (mg), and relative TA muscle mass (mg/g of body weight) of uninjured young (4-6 mo) and aged (26-28 mo) female C57BL/6 mice. B: Absolute maximal nerve stimulated *in-situ* isometric contractile force (P_o_, mN) generated by young and aged TA muscles was measured and used to calculate maximal specific isometric contractile force (sP_o_, mN/mm^2^). Representative force traces obtained from young and aged mice are shown. C: TA muscle cross-sections were stained with antibodies for type I, IIa, and IIb myosin heavy chain and stitched panoramic 4-color fluorescent images of the entire muscle cross-section obtained. Type IIx fibers remain unstained (black). The extracellular matrix was labeled with wheat germ agglutinin (WGA) to delineate myofiber borders. Scale bars are 1000 μm. Right panels show representative images of single fields of view obtained from deep and superficial regions of the TA showing a clear influence of anatomical location on both myofiber size and fiber type. Scale bars are 200 μm. Therefore, the total number and cross-sectional area (CSA) of each muscle fiber and its corresponding fiber type throughout the entire muscle cross-section was determined via high throughput fully automated image analysis using the MuscleJ plugin for FIJI/ImageJ. D: Quantification of the total number of muscle fibers, mean myofiber cross-sectional area (CSA), and frequency distribution histogram of myofiber CSA as determined by MuscleJ. E: Quantification of TA muscle fiber type composition and mean fiber CSA split by respective fiber type as determined by MuscleJ. Bars show the mean ± SEM of 5-7 mice per group with dots representing data from each individual mouse. *Denotes p<0.05 vs. young mice by a two-tailed unpaired t-test.

### Chronic Basal Inflammation of Aged Muscle is Associated with a Lack of Anti-Inflammatory and Pro-Resolving Lipid Mediators

In order to investigate the mechanisms that may contribute to age-associated muscle atrophy and weakness, we first performed LC-MS based metabolipidomic profiling of uninjured TA muscle samples from young and aged mice. Unsupervised principle component analysis (PCA) showed a clear clustering of samples by age (Figure 2A). The corresponding loadings of some key representative metabolites from each major enzymatic pathway that contributed to this response are shown in Figure 2B. The lipid mediator profile of young muscle was characterized by an abundance of a range of bioactive PUFA metabolites, many of which were derived from the 5-LOX and 15-LOX pathways. Of the 98 total lipid mediator species detected in muscle tissue, 59 were present at >1.5-fold lower concentration in aged vs young muscle (Figure 2C). In contrast, only a single analyte tended to be enriched in aged muscle (Figure 2C). Comparison of lipid mediator abundance pooled by respective enzymatic pathway revealed that total COX-derived prostaglandins were similarly abundant in young and aged muscle (Figure 2D, Supplemental Table 1A). In contrast, monohydroxy-fatty acids derived from the 5- and 15-LOX (and to a lesser extent 12-LOX) pathways were substantially lower in aged mice (Figure 2D, Supplemental Table 1B). Mature SPMs, which are produced endogenously by the sequential action of two or more LOX isoforms, were present at very low concentrations in basal muscle and were often below the limits of detection (Supplemental Table 1D). Nevertheless, lipoxin A_4_ (LXA_4_), resolvin D6 (RvD6), and maresin 1_n-3 DPA_ (MaR1_n-3DPA_) were detected in all samples, while resolvin E3 (RvE3), maresin 1 (MaR1), and 8-oxo-RvD1 were detected only in young muscle (Figure 2C, Supplemental Table 1D). This resulted in an overall basal deficiency of pooled SPMs in aged mice (Figure 2D). The activity of the cytochrome p450 (CYP) pathway, which primarily generates fatty acid epoxides [e.g. epoxyeicosatrienoic acids (EpETrEs)] with potent anti-inflammatory actions was also markedly lower in aged muscle prior to injury (Figure 2D, Supplemental Table 1C).

**Figure 2.**
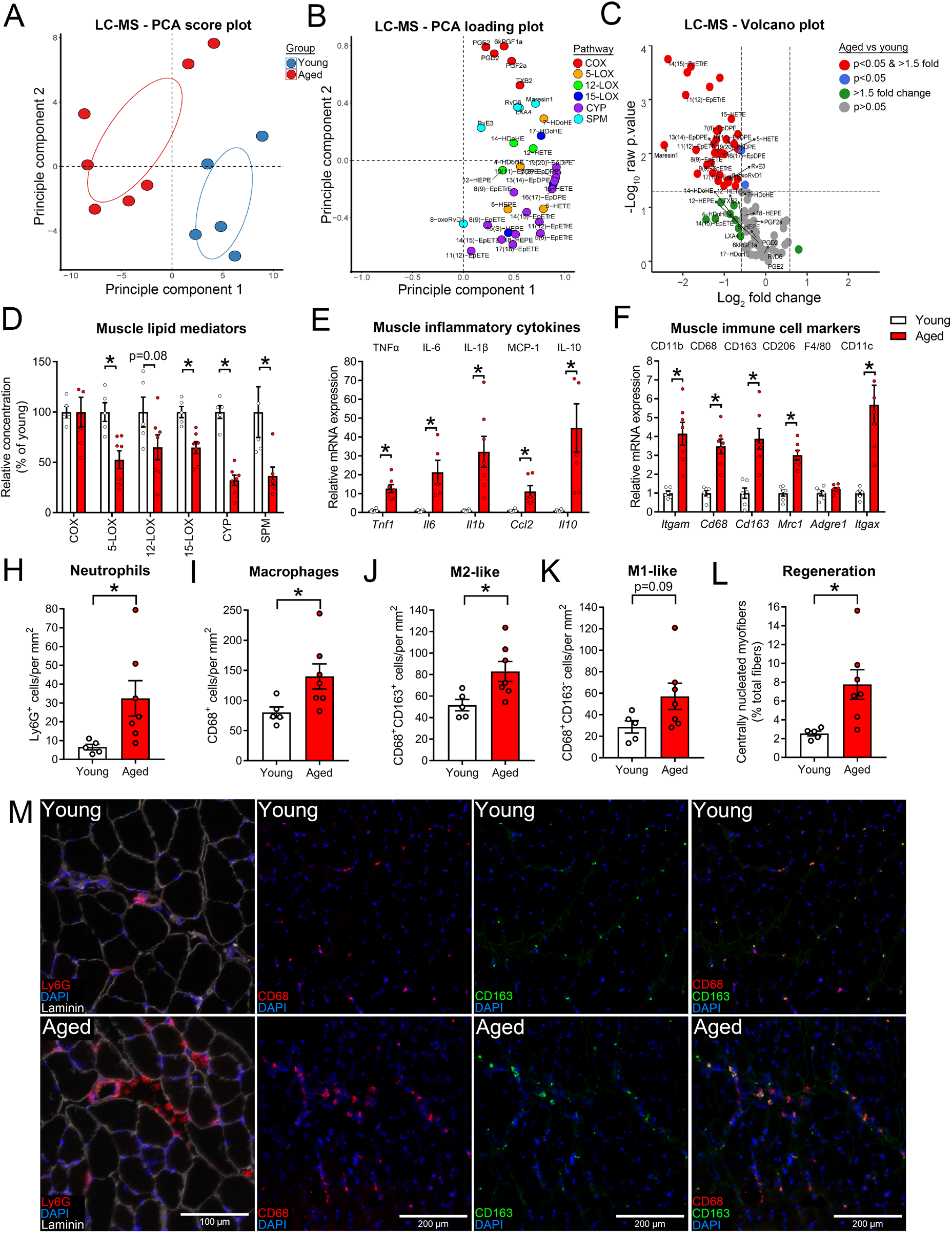
Chronic low-grade inflammation of aged skeletal muscle is associated with a basal deficiency of anti-inflammatory and pro-resolving lipid mediators: A-B: Unsupervised principle component analysis (PCA) score and loading plots of the entire mediator lipidome (143 analytes) of uninjured tibialis anterior (TA) muscles from young (4-6 mo) and aged (26-28 mo) female C57BL/6 mice as determined by liquid chromatography-tandem mass spectrometry (LC-MS/MS)-based metabolipidomic profiling. C: Volcano plot displaying the magnitude of difference between aged vs young mice and related statistical significance (as unadjusted p-values) for each individual detected lipid mediator (98 analytes total) D: Relative pooled concentrations of muscle lipid mediators derived from each major enzymatic biosynthesis pathway in young and aged mice. E-F: Whole TA muscle mRNA expression of inflammation-related genes including inflammatory cytokines and immune cell surface markers and as determined by real-time quantitative reverse transcription PCR (RT-qPCR). H-L: Quantification of immunohistochemical staining of TA muscle cross-sections for intramuscular number of polymorphonuclear cells (PMNs) (Ly6G^+^ cells), total macrophages (MΦ) (CD68^+^ cells), M2-like MΦ (CD68^+^CD163^+^ cells), M1-like MΦ (CD68^+^CD163^−^ cells), and degenerating/regenerating (centrally nucleated) myofibers. M: Representative staining of myeloid cell populations in cross-sections of young and aged TA muscle samples. Scale bars are 200 μm. Bars show the mean ± SEM of 5-7 mice per group with dots representing data from each individual mouse. *Denotes p<0.05 vs vehicle group by a two-tailed unpaired t-test.

Age-associated deregulation of the skeletal muscle mediator lipidome was accompanied by heightened whole muscle mRNA expression of pro-inflammatory cytokines and immune cell markers (Figure 2E-F), as well as greater intramuscular numbers of PMNs, total MΦ, M2-like MΦ, and to a lesser extent M1-like MΦ as determined by immunohistochemistry (Figure 2H-K). This accumulation of predominantly resident M2-like MΦ in aged muscle was further confirmed by quantitative flow cytometry of the total single-cell population isolated from whole hind-limb muscles from young and aged mice (Supplemental Figure 1). Chronic inflammation of aged muscle was also related to ongoing myofiber degeneration/regeneration as evidenced by an increased presence of centrally nucleated muscle fibers (Figure 2L). Representative images of immunohistochemical staining of intramuscular myeloid cell populations in young and aged muscle tissue are shown in Figure 2M. Overall these data show that chronic inflammation of aging skeletal muscle is characterized by a basal deficiency of many anti-inflammatory and pro-resolving lipid mediators derived from both the LOX and CYP pathways.

### Aged Mice Mount a Deficient Specialized Pro-Resolving Lipid Mediator Response to Muscle Injury

We next assessed the effect of aging on the bioactive lipid metabolome of muscle following sterile injury induced by intramuscular injection of barium chloride (BaCl_2_). There was a minor shift in global muscle lipid mediator profile at day 1 post-injury in young mice (Figure 3A). This was predominantly due to a rapid increase in likely non-enzymatic PUFA metabolites (Figure 3B, Supplemental Table 1E). This was followed by a marked shift in muscle lipid mediator profile at day 3 post-injury attributable to increased concentrations of many major enzymatic products of the COX, LOX, and CYP pathways (Figure 3A-B, Supplemental Tables 1A-D). There was a further transition from day 3 to day 5 of recovery from muscle injury from an earlier predominance of 5-LOX and CYP metabolites towards to a more delayed and prolonged elevation of COX and 12-LOX pathway products.

**Figure 3.**
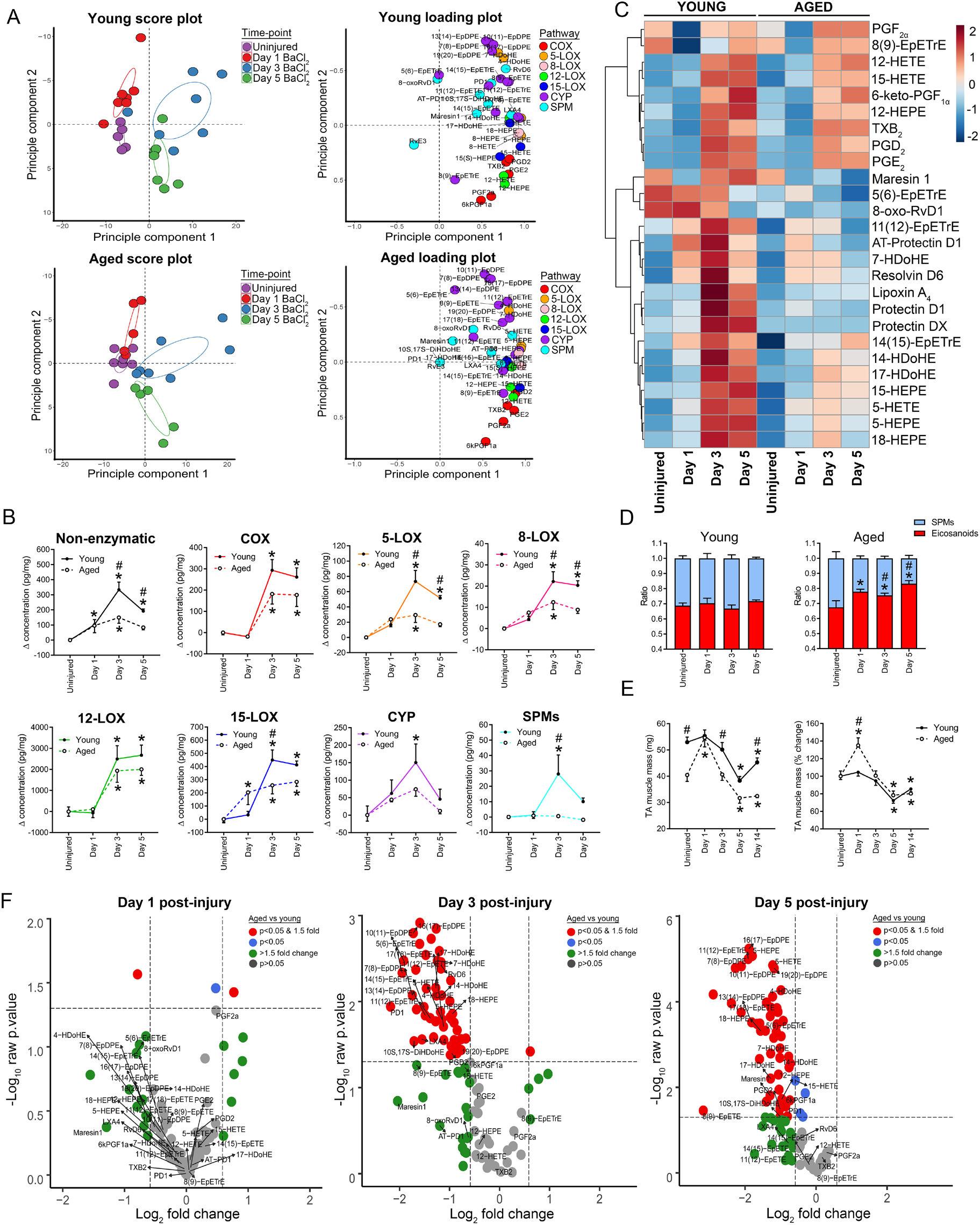
Lipidomic profiling of the impact of age on skeletal muscle injury and repair: A: Young (4-6 mo) and aged (26-28 mo) female C57BL/6 mice received bilateral intramuscular injection of the tibialis anterior (TA) muscle with 50 μL of 1.2% barium chloride (BaCl_2_) to induce myofiber injury. TA muscles were collected at 1, 3, and 5 days’ post-injury for analysis of intramuscular lipid mediator concentrations via LC-MS/MS based metabolipidomic profiling. Principle component analysis (PCA) score and loading plots displaying global shifts in the mediator lipidomic profile of injured muscle over the time-course of inflammation-resolution. B: Pooled concentrations of lipid mediators from various major potential biosynthetic routes following muscle injury. Data is baseline corrected using the average resting concentration of lipid mediators in age-matched uninjured young and aged muscle as displayed in Figure 2D (the Δ change). C: Heat map and hierarchical clustering of temporal shifts in major individual representative lipid mediators from the cyclooxygenase (COX), lipoxygenase (LOX), and epoxygenase (CYP) pathways. D: Shifts in the balance of pro-inflammatory eicosanoids relative to the specialized pro-resolving mediators (SPMs) and their related monohydroxy-PUFA pathway markers (15-HETE, 14-HDoHE, 17-HDoHE, 18-HEPE) following muscle injury. E: Absolute and relative changes in the total mass of the TA muscle samples used for lipidomic profiling. F: Volcano plots displaying the magnitude and related statistical significance (as unadjusted p-values) between aged and young mice for each individual detected analyte at day 1, 3, and 5 following muscle injury. Bars show the mean ± SEM of 5-7 mice per group. *denotes p<0.05 change over time when compared to uninjured muscles as determined by two-way ANOVA with Holm-Sidak post-hoc tests. #denotes p<0.05 difference between young and aged mice at a given time-point by two-way ANOVA with Holm-Sidak post-hoc tests.

When compared to young mice, aged mice mounted an overall deregulated lipid mediator response as evidenced by less separation of clusters of samples obtained at distinct time-points (Figure 3A). Parametric statistical analysis of intramuscular lipid mediators pooled over major biosynthetic pathways (by two-way ANOVA) revealed that aged and young mice produced similar amounts of COX, 12-LOX, and CYP products following injury (age × time p=0.18, p=0.66, p=0.64 respectively) (Figure 3B). In contrast, aged mice showed diminished local production of 5-, 8- and 15-LOX metabolites, as well as a marked deficiency in downstream bioactive SPMs (age × time effects of p=0.0017, p=0.018, p=0.0053, p=0.011 respectively) (Figure 3B). Non-enzymatic lipid mediator production was also lower in aged mice in response to muscle injury (age × time p=0.0072) (Figure 3B).

A heat map displaying temporal shifts in key representative individual lipid mediator species from each of these major biosynthetic pathways following muscle injury is shown in Figure 3C. The age-related deficiency of biosynthesis of SPMs including LXA_4_, RvD6, PD1, 10S,17S-DiHDoHE [also known as protectin DX (PDX)], MaR1, MaR1_n-3DPA_, 8-oxo-RvD1 (Supplemental Table 1D), as well as their key pathway markers/intermediates (e.g. 15-HETE, 14-HDoHE, 17-HDoHE, 18-HEPE) (Supplemental Table 1B), when combined with a largely equivalent production of major pro-inflammatory COX (e.g. PGE_2_) and 12-LOX (e.g. 12-HETE) metabolites (Supplemental Table 1A), resulted in a relative overabundance of classical eicosanoids as evidenced by the SPM:eicosanoid ratio (Figure 3D). With the exception of an initial transient increase in muscle weight at day 1 post-injury in aged mice only, changes in the total mass of the TA muscles used for lipidomic profiling over the time-course of recovery from muscle injury was similar between young and aged mice and thus did not appear to contribute substantially to the effect of age on muscle lipid mediator concentration (Figure 3E). Volcano plots summarizing the overall differences between aged and young muscle mediator lipidomes for all analytes detected by LC-MS/MS are shown in Figure 3F. Overall these findings show that aging results in a marked imbalance in local biosynthesis of pro-inflammatory and anti-inflammatory/pro-resolving lipid mediators following sterile muscle injury.

### SPM Treatment Suppresses Intramuscular Inflammatory Cytokines, but Does Not Limit Excessive Leukocyte Infiltration of Aged Muscle

Since aging was associated with a markedly impaired SPM response to muscle injury, we tested whether exogenous SPM treatment could potentially influence muscle inflammation following injury in aged mice. We first confirmed the established bioactivity of the prototypical SPM resolvin D1 (RvD1) on bone marrow derived macrophages (BMM) *in-vitro.* RvD1 potently stimulated phagocytosis by BMM obtained from both young mice and aged mice (Supplemental Figure 2). These data confirm the previously reported bioactivity of RvD1 on myeloid cells in our hands and demonstrate that aged MΦ appear to maintain the intrinsic ability to respond effectively to RvD1.

We then treated aged mice *in-vivo* with RvD1 by daily intraperitoneal injection following muscle injury with the first dose administered approximately 5 min prior to intramuscular BaCl_2_ injection. There was a large increase in intramuscular presence of PMNs (Ly6G^+^ cells) and MΦ (CD68^+^ cells) at day 1 post-injury (Figure 4A). This was followed by a subsequent decline in PMN number and progressive MΦ recruitment peaking at day 3 of recovery (Figure 4B). When compared to young mice, aged mice initially showed relatively greater recruitment of both PMNs and MΦ at day 1 post-injury (Figure 4C). However, by day 3 post-injury aged mice showed more rapid PMN clearance together with a greater intramuscular MΦ presence (Figure 4D). Treatment of aged mice with RvD1 did not influence the number of PMNs or MΦ in muscle as determined by immunohistochemical staining of tissue cross-sections between day 1 and 3 post-injury (Figure 4C-D).

**Figure 4.**
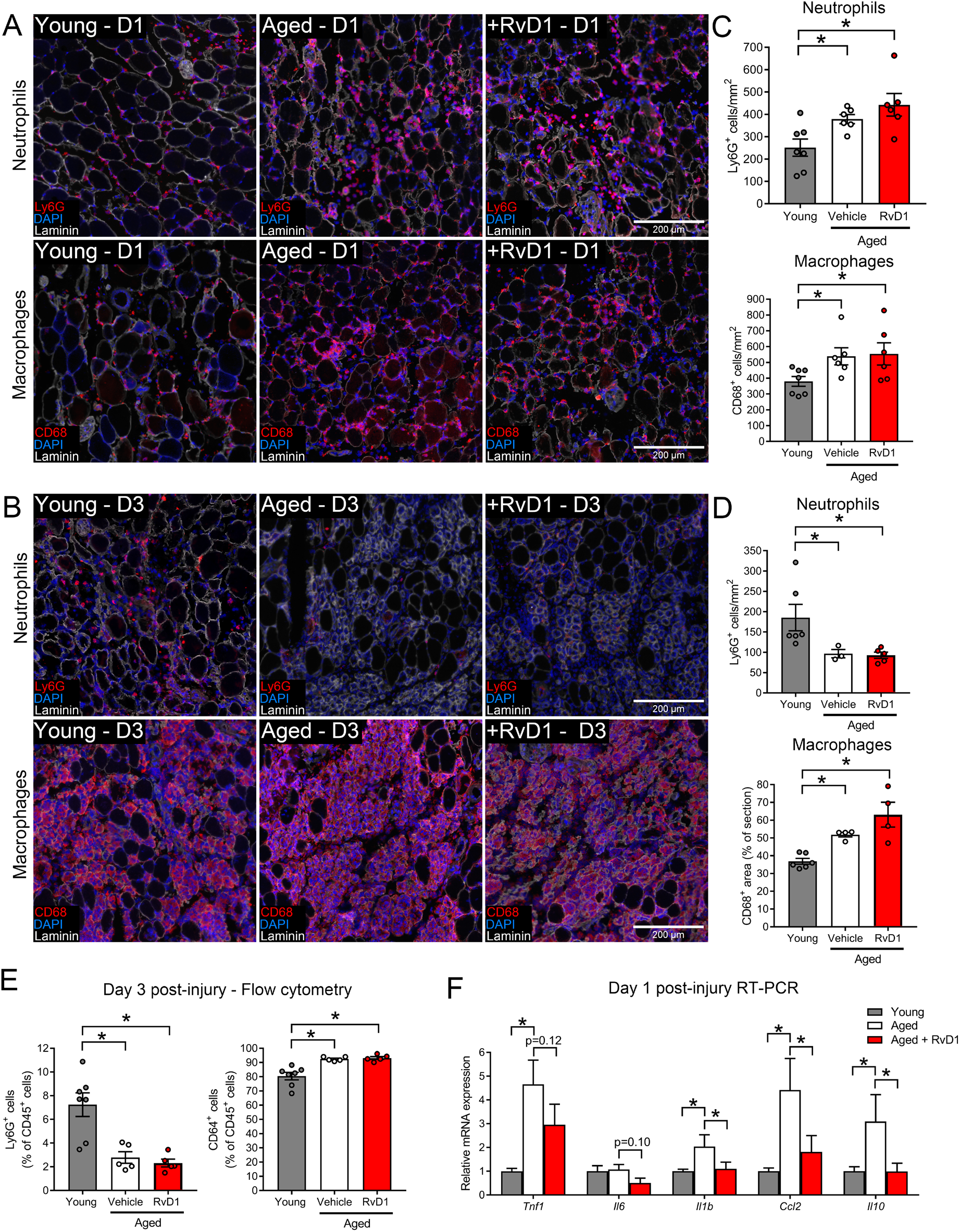
The effect of aging and resolvin D1 treatment on acute myeloid cell responses to skeletal muscle injury: A-B: Young (4-6 mo) and aged (26-28 mo) female C57BL/6 mice received bilateral intramuscular injection of the tibialis anterior (TA) muscle with 50 μL of 1.2% barium chloride (BaCl_2_) to induce myofiber injury. Aged mice received daily treatment with resolvin D1 (RvD1, 100 ng) or vehicle (0.1% ethanol) via intraperitoneal (IP) injection with the first dose administered ~5 min prior to muscle injury. TA muscles were collected at day 1 (D1) and day 3 (D3) post-injury. Muscle cross-sections were stained for either polymorphonuclear cells (PMNs, Ly6G) or monocytes/macrophages (MΦ, CD68). Cell nuclei and the basal lamina were counterstained with DAPI and a laminin antibody respectively. Scale bars are 200 μm. C: Quantification of the number of PMNs (Ly6G^+^ cells) and MΦ (CD68^+^ cells) (both as cell counts) in injured muscle at day 1 post-injury. D: Quantification of the number of PMN (Ly6G^+^ as cell counts) and total MΦ^+^ staining area (as % of total tissue) in injured muscle at day 3 post-injury. E: Single-cells isolated from pooled left and right TA muscles of young and aged mice at day 3 post-injury were analyzed by flow cytometry with antibodies for CD45-PE, Ly6G-FITC, and CD64-APC. Dead cells were excluded from analysis using LIVE/DEAD fixable violet. Intramuscular PMNs (Ly6G^+^ cells) and MΦ (CD64^+^ cells) were quantified as percentage of total intramuscular leukocytes (CD45^+^ cells). F: Quantification of whole muscle mRNA expression of inflammatory cytokines at day 1 post-injury as determined by RT-qPCR. Bars show the mean ± SEM of 4-7 mice per group with dots representing data from each individual mouse. *Denotes p<0.05 between groups by one-way ANOVA with pairwise LSD post-hoc tests.

To confirm the surprising finding that despite their markedly deficient local SPM response that aging actually appeared to increase the rate of clearance of PMNs from injured muscle, we repeated these experiments for analysis of the entire intramuscular single-cell population by flow cytometry. At day 3 post-injury ~8% of intramuscular leukocytes (CD45^+^ cells) were PMNs (Ly6G^+^ cells) in young mice, compared with only ~2% in aged mice (Figure 4E). This was accompanied by a parallel increase in the relative proportion of MΦ (CD64^+^ cells) in aged muscle. Treatment of aged mice with RvD1 did not influence the proportion of CD45^+^ cells that were either PMN or MΦ in injured muscle (Figure 4E). Nevertheless, RvD1 treatment did reduce local mRNA expression of inflammatory cytokines within injured muscle tissue as determined by RT-PCR (Figure 4F). These data show that RvD1 may still influence the inflammatory profile of intramuscular myeloid cells in aged mice despite apparently not impacting their presence.

### Defective Myofiber Regeneration in Aged Mice is not Influenced by RvD1

At day 5 post-injury, there was an appearance of many small regenerating myofibers with characteristic centrally located nuclei that robustly expressed embryonic myosin heavy chain (eMHC) in both young and aged mice (Figure 5A). When compared to young mice, regenerating muscle cross-sections from aged mice were overall smaller in size, contained fewer total muscle fibers, and had far less newly formed myofibers (as assessed by the combination of eMHC expression and associated centrally located myonuclei) (Figure 5B). In addition to a lack of fibers undergoing regeneration in aged mice, the myofibers that were regenerating were far smaller in size (Figure 5C-D). Daily treatment of aged mice with RvD1 did not influence overall muscle size, total myofiber number, regenerating fiber number, or the average size of the regenerating myofiber population (Figure 5A-D).

**Figure 5.**
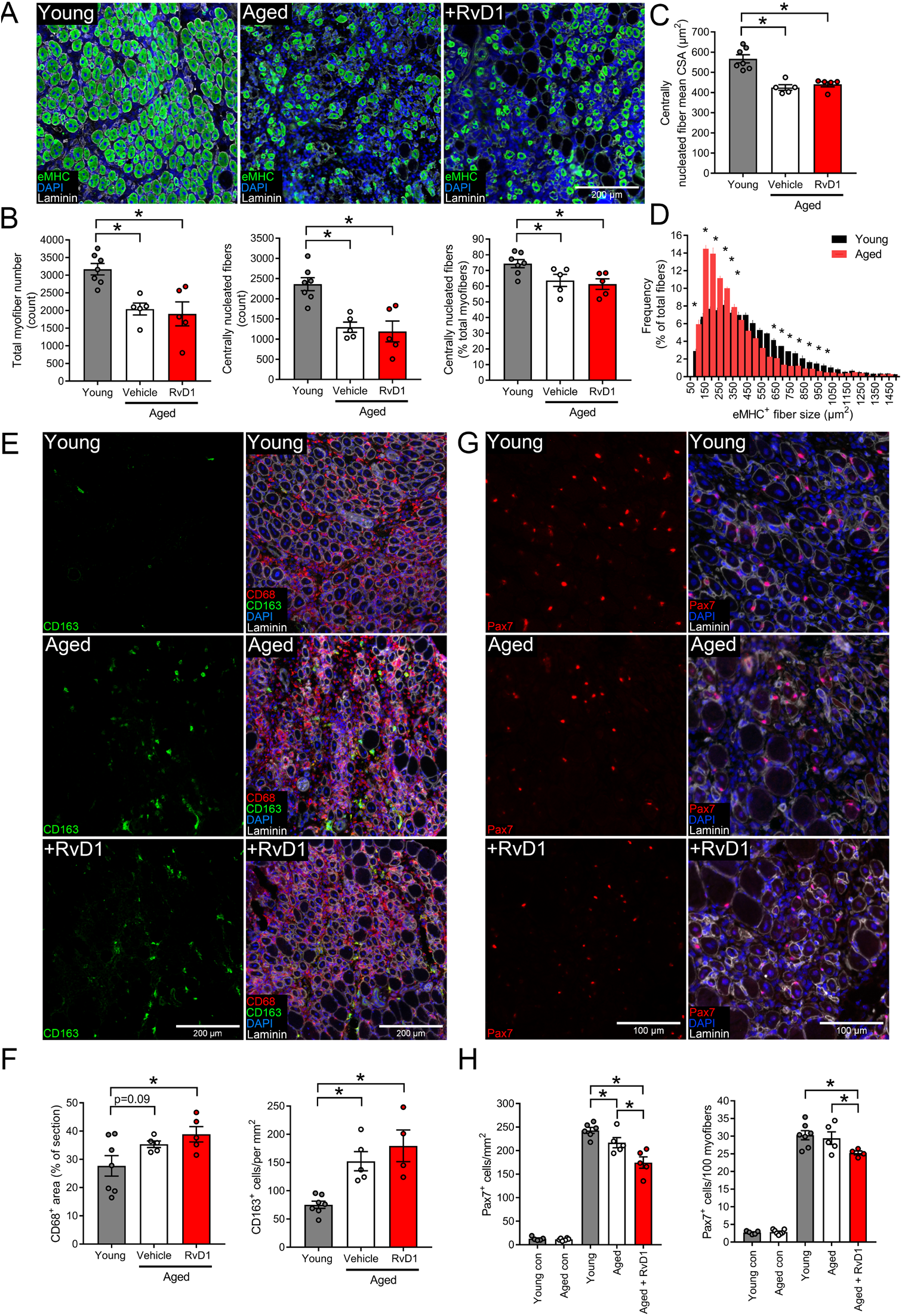
The effect of age and resolvin D1 treatment on myofiber regeneration: A: Young (4-6 mo) and aged (26-28 mo) female C57BL/6 mice received bilateral intramuscular injection of the tibialis anterior (TA) muscle with 50 μL of 1.2% barium chloride (BaCl_2_) to induce myofiber injury. Aged mice were then treated daily with resolvin D1 (RvD1) (100 ng) or vehicle (0.1% ethanol) by intraperitoneal (IP) injection with the first dose administered ~5 min prior to injury. TA muscles were collected at day 5 post-injury and muscle cross-sections were stained for embryonic myosin heavy chain (eMHC) to label newly formed myofibers. Cell nuclei and the basal lamina were counterstained with DAPI and a laminin antibody respectively. Scale bars are 200 μm. B: Quantitative analysis of total myofiber number, regenerating (centrally nucleated) myofiber number, and percentage of total myofibers undergoing regeneration (≥1 central nuclei) as determined by MuscleJ. C: The mean cross-sectional area (CSA) of the regenerating myofiber population as determined by MuscleJ. D: Frequency distribution histogram of regenerating TA myofiber CSA in young and aged mice as determined by MuscleJ. E: Regenerating muscle cross-sections were stained for total MΦ (CD68) and M2-like MΦ (CD163). Scale bars are 200 μm. F: Total MΦ staining was determined as CD68^+^ tissue area (mm^2^) and expressed relative to total muscle cross-section CSA (%) as determined by MuscleJ. M2-like MΦ (CD163 cells) were manually counted throughout the entire section and as expressed relative to total muscle CSA as determined by MuscleJ. G: Regenerating muscle cross-sections were stained for the muscle satellite cell (MuSC) marker Pax7. TA muscles from age and gender matched uninjured mice were analyzed to determine the effect of aging on resting MuSC number. Scale bars are 100 μm. H: Quantification of the number of MuSCs (Pax7^+^DAPI^+^ nuclei located beneath the basal lamina) expressed relative to tissues area (per mm^2^) or myofiber number (per 100 fibers). Bars are mean ± SEM of 5-7 mice per group with dots representing data for each individual mouse. *p<0.05 by one-way ANOVA with pairwise LSD post-hoc tests for panels A-F or pairwise Holm-Sidak post-hoc tests for panel H.

When compared to young mice, regenerating aged muscles tended to have more total MΦ at day 5 post-injury (p=0.09), and this was especially true for M2-like MΦ (CD68^+^CD163^+^ cells) (Figure 5E). However, treatment of aged mice with RvD1 did not impact intramuscular MΦ numbers (Figure 5F). Consistent with their important role in muscle regeneration, muscle satellite cell (MuSC) number also increased markedly at day 5 post-injury when compared to uninjured muscles (Figure 5G). Aged mice showed lower numbers of MuSCs at this time-point than young mice and this was reduced even further in aged mice treated with RvD1 (Figure 5H). Collectively this data shows that RvD1 treatment does appear to have a marked impact on the MuSC response to muscle damage, but that this does not appear to translate to a clear positive or negative impact on myofiber regeneration in aged muscle.

### Resolvin D1 Limits Maladaptive Remodeling of Aged Muscle and Improves Recovery of Muscle Function

When expressed as percent deficit compared to the average strength of age-matched uninjured muscles (as shown in Figure 1), aged injured mice had greater functional deficits than young injured mice for maximal isometric contractile force (P_o_) and much of this difference persisted after accounting for their smaller regenerating muscle size (sP_o_) (Figure 6A). In contrast, aged mice treated with RvD1 did not significantly differ from young mice for deficits in either P_o_ or sP_o_. Moreover, RvD1 treatment tended to result in an improvement in the sP_o_ deficit when compared to aged mice receiving vehicle treatment (p=0.06). This finding indicated that the benefit of RvD1 treatment on normalized force generating capacity (sP_o_) appeared to be driven by a parallel decrease in whole muscle cross-sectional area.

**Figure 6.**
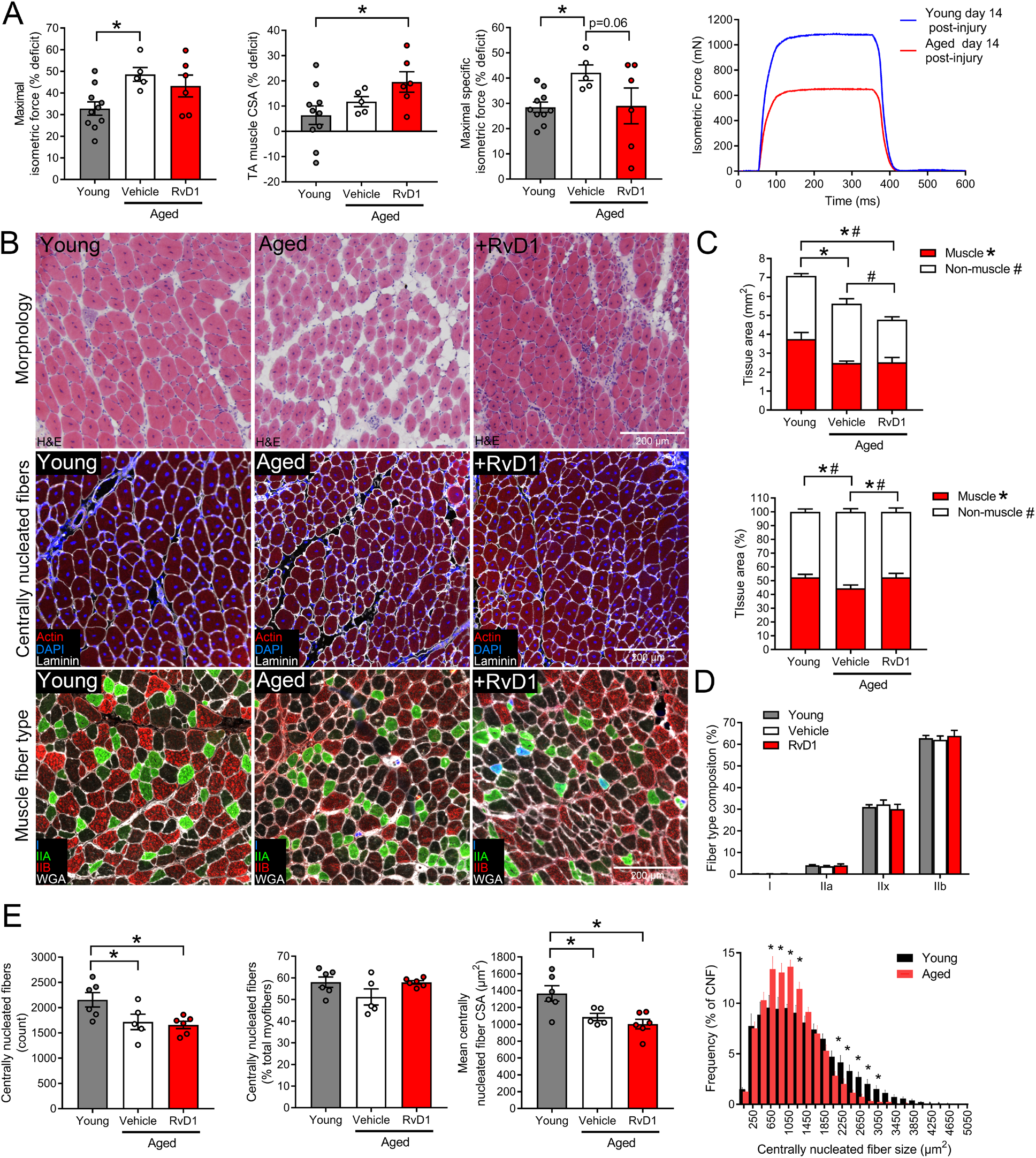
Resolvin D1 limits maladaptive tissue remodeling following skeletal muscle injury in aged mice: A: Young (4-6 mo) and aged (26-28 mo) female C57BL/6 mice received bilateral intramuscular injection of the tibialis anterior (TA) muscle with 50 μL of 1.2% barium chloride (BaCl_2_) to induce myofiber injury. Aged mice were then treated with daily intraperitoneal (IP) injections of resolvin D1 (RvD1) (100 ng) or vehicle (0.1% ethanol) for 14 days with the first dose administered ~5 min prior to injury. At two-weeks post-injury (day 14), absolute maximal nerve-stimulated *in-situ* isometric contractile force (P_o_, mN) generated by the TA muscle was measured and used to calculate specific force (sP_o_, mN/mm^2^). Data is presented as percentage force deficit relative to the strength of age-matched uninjured TA muscles shown in Figure 1. B: Day 14 post-injury TA cross-sections were stained for hematoxylin & eosin (H & E), conjugated phalloidin to label all muscle fibers (actin), or with antibodies for type I, IIa, and IIb myosin heavy chain to label specific muscle fiber types. Type IIx fibers remain unstained (black). On actin slides cell nuclei and the basal lamina were counterstained with DAPI and a laminin antibody respectively to identify centrally nucleated (regenerating) myofibers. On fiber type slides the extracellular matrix was stained with wheat germ agglutinin (WGA) to delineate myofiber boundaries. Scale bars are 200 μm. All image analysis was performed by fully automated analysis of the entire muscle cross-section using the MuscleJ plugin for FIJI. C: Total cross-sectional area of regenerating TA muscles and relative amounts of tissue area containing myofibers (actin^+^ tissue area) compared with that devoid of myofibers (actin^−^ tissue area). D: Quantification of the fiber type composition of regenerating muscles from young and aged mice. E: Quantification of total regenerating (centrally nucleated) myofiber number, the percentage of total fibers undergoing regeneration, and the mean cross-sectional area (CSA) of the regenerating myofiber population. Frequency distribution histograms of regenerating myofiber CSA for young vs vehicle treated aged mice are shown. Bars show mean ± SEM of 5-10 mice per group with dots representing data from each individual mouse *Denotes p<0.05 by one-way ANOVA with pairwise LSD post-hoc tests.

To investigate this further, we first performed H & E staining and showed that compared to young mice, regenerating muscles from untreated aged mice had a greatly expanded interstitial space between their smaller regenerating myofibers (Figure 6B). Cross-sections of aged TA muscles were overall smaller in size than young mice and this was mainly due to a reduction in functional muscle fiber (actin^+^) area, while the total amount of non-myofiber (actin^−^) tissue (e.g. extracellular matrix) was similar to young muscle (Figure 6C). Because of this, regenerating aged muscles were comprised of a relatively greater proportion of non-contractile fibrotic and/or fatty tissues, indicative of poorer muscle quality. When compared to aged mice receiving vehicle treatment, RvD1 did not influence overall muscle cross-sectional area or the absolute amount of myofiber (actin^+^) containing area (Figure 6C). But, RvD1 treatment did significantly reduce the accumulation of fibrotic/fatty (actin^−^) tissues within regenerating muscle (Figure 6C). Consequently, the relative proportion of regenerating muscle which was compromised of functional contractile muscle fibers was increased in aged mice treated with RvD1 when compared to aged mice not receiving treatment.

Neither age nor RvD1 treatment had a significant impact on the fiber type composition of regenerating muscles (Figure 6D) and both young and aged TA muscles were predominantly composed of regenerating (centrally nucleated) myofibers at day 14 post-injury (Figure 6E). When compared to young mice, aged muscles contained relatively fewer regenerating myofibers which were on average smaller in size and this too was not influenced by RvD1 treatment. Aged muscles also still contained many more intramuscular MΦ than young mice even at 14 days following muscle damage and treatment of aged mice with RvD1 did not significantly influence this response (Supplemental Figure 3).

## Discussion

Here we examined the effect of aging on the local lipid mediator response to muscle injury and tested whether systemic RvD1 treatment could limit excessive inflammation and improve muscle regeneration in aged mice. When compared to young muscle, there was increased basal inflammatory cytokines and accumulation of both PMNs and M2-like MΦ in aged muscle. This chronic low-grade inflammation was related to a deficiency in anti-inflammatory and pro-resolving lipid mediators within muscle even prior to injury. Following injury, aged mice displayed a markedly deficient intramuscular SPM response, which was associated with excessive acute muscle inflammation, deficient myofiber regeneration, and impaired recovery of muscle strength. Treatment of aged mice with RvD1 limited the local inflammatory cytokine response, but did not impact myeloid cell recruitment or myofiber regeneration. Nevertheless, RvD1 did modulate the MuSC response, limited maladaptive tissue remodeling, and improved recovery of relative muscle function. These findings suggest that age-associated muscle dysfunction is related to a striking imbalance of intramuscular pro-inflammatory and anti-inflammatory/pro-resolving lipid mediators. Further, immunoresolvents may be an attractive alternative for the clinical management of muscular injuries and associated pain in the elderly, due to positive effects on muscle quality and functional recovery of muscle strength without the negative side effects on myofiber regeneration of traditional non-steroidal anti-inflammatory drug treatments.

Consistent with recent studies, we found that when compared to young mice, aged muscle showed an increased basal presence of both PMNs (Sloboda et al., 2018) and resident M2-like MΦ (Cui et al., 2019; Reidy et al., 2019; Sorensen et al., 2019; Wang et al., 2015). Aged mice also mounted a heightened acute inflammatory response to myofiber injury induced by intramuscular injection of BaCl_2_ which is consistent with prior studies of muscle damage induced by lengthening contractions (Koh, Peterson, Pizza, & Brooks, 2003; Sloboda et al., 2018), myotoxin injection (Patsalos et al., 2018; van der Poel et al., 2011), or contusion (Ghaly & Marsh, 2010). LC-MS/MS based profiling of the mediator lipidome revealed that both chronic low-grade muscle inflammation and excessive acute myeloid cell responses in muscle injury were associated with a marked deficiency in local concentrations of SPMs and their key pathway markers, suggesting that pro-resolving therapies may benefit age-related muscle dysfunction.

Treatment of aged mice with the prototypical SPM, RvD1, at the time of injury suppressed muscle expression of both pro- and anti-inflammatory cytokines including MCP-1, IL-1β, and IL-10. Despite this, unlike our prior study in young mice (Markworth et al., 2020), in the present study we were unable to demonstrate in aged mice a reduction in MΦ infiltration of muscle in response to RvD1 treatment. Moreover, other inflammation-related genes (e.g. TNFα and IL-6) that were markedly suppressed in young mice treated with RvD1 (Markworth et al., 2020), showed only statistical trends towards suppression in aged muscle here. These data suggest that RvD1 was unable to fully overcome heightened muscle myeloid cell infiltration in aged mice. This may relate to the fact that as in prior studies expression of many inflammation-related genes was increased in aged muscle even before injury (Cui et al., 2019; Lavin et al., 2020; Patsalos et al., 2018; Rivas et al., 2016; van der Poel et al., 2011; Wang et al., 2015; Wang et al., 2018). Furthermore, the degeneracy of SPMs may also suggest that a single immunoresolvent class (RvD1 used here) may be insufficient for the overall resolution of inflammation following muscle injury, especially in aged mice that were deficient in a range of different SPM families. Therefore, future studies should examine whether longer-term immunoresolvent or a combinatorial SPM treatment may reduce chronic basal inflammation of aged skeletal muscle and the effect this may have on the inflammatory and adaptive responses to subsequent muscle damage.

We further observed an unanticipated and marked deficit in many fatty acid epoxides derived from the third and less well explored CYP lipid mediator pathway in aged muscle prior to injury. These bioactive oxylipins which include the four major epoxyeicosatrienoic acid regioisomers derived from n-6 arachidonic acid [5,6-, 8,9, 11,12, and 14,15-EpETrE (also known as EETs)] as well as analogous metabolites of n-3 EPA (EpETEs) and DHA (EpDPEs) possess potent anti-inflammatory actions (Christmas, 2015). On this basis, their reduced presence within aged muscle may also contribute to basal inflammation in addition to a lack of LOX metabolites (e.g. SPMs, HEPEs, HDoHEs). A recent study showed that CYP metabolites also display similar actions to the SPMs in stimulating resolution of the acute inflammatory response (Gilroy et al., 2016). Moreover, the CYP pathway contributes to endogenous biosynthesis of the E-series resolvins (e.g. RvE1) by producing the key pathway marker and intermediate 18-HEPE (Arita, Clish, & Serhan, 2005). Therefore, metabolites of the LOX and CYP pathways are likely to act in unison. While we focused our intervention on SPMs here, as biosynthesis of these particular LOX metabolites were most lacking in response to sterile muscle injury, our data also suggest that targeting the CYP pathway may also be a novel and previously unexplored strategy to combat basal age-associated muscle inflammation and related dysfunction. One such strategy is to block conversion of bioactive anti-inflammatory lipid epoxides to their inactive downstream vicinal diols [DHETrEs (also known as DHETs)] via inhibition of soluble epoxide hydrolase (sEH) (Wagner, McReynolds, Schmidt, & Hammock, 2017).

The ability of systemic RvD1 administration to limit PMN recruitment is well-described for certain experimental models of acute inflammation (Norling, Dalli, Flower, Serhan, & Perretti, 2012). In contrast, we have been unable to demonstrate this in the context of sterile skeletal muscle injury in either young (Markworth et al., 2020), or old mice (present study). Prior studies in cardiac muscle found that systemic injection of RvD1 was also unable to limit PMN appearance following myocardial infarction (Halade, Kain, & Serhan, 2018; Kain et al., 2015). Therefore, the suppressive effects of SPMs on PMN recruitment may depend both on the nature of the inflammatory insult as well as the site of inflammation. SPMs such as RvD1 also enhance PMN clearance via distinct mechanisms of action, thereby accelerating a return to a non-inflamed state (Bannenberg et al., 2005). Indeed, repeated daily administration of RvD1 reduced intramuscular PMN numbers by day 3 of recovery from muscle injury in young mice in our prior study (Markworth et al., 2020). Therefore, we hypothesized that RvD1 treatment would hasten clearance of the greater numbers of PMNs, which invaded aged muscle early following injury. To our surprise, aged mice actually cleared muscle PMNs much more rapidly than young mice and RvD1 treatment was unable to further accelerate this response. PMNs can inflict secondary muscle damage and limiting the influx of intramuscular PMN response is generally considered to be organ protective in the context of muscle injury (Pizza, Peterson, Baas, & Koh, 2005). In this context, it is surprising that aging resulted in both an enhanced rate of PMN clearance from muscle as well as defective myofiber regeneration. While evidence from the skeletal muscle literature is currently lacking, PMNs do also appear play an important role in tissue repair under certain physiological conditions (Peiseler & Kubes, 2019). For example, PMNs have been shown to support dermal wound healing in aged mice (Nishio, Okawa, Sakurai, & Isobe, 2008). Given the marked effect of aging on the increased magnitude but decreased duration of the PMN response to muscle injury identified in the current study, future studies are needed to better clarify the role of PMNs in age-related muscle dysfunction.

The lack of PMNs within aged muscle during the resolution phase of the inflammatory response was accompanied by greater number of tissue MΦ. These data are consistent with prior studies of both myotoxin injection (Patsalos et al., 2018) and contusion injury (Ghaly & Marsh, 2010). However, these findings conflict with recent studies of more mild muscle inflammation induced by either recovery from disuse (Reidy et al., 2019) or contraction-induced injury (Sloboda et al., 2018), in which lower numbers of MΦ were observed in aged muscle. The role of the greater local MΦ accumulation in aged compared with young mice following degenerative muscle injury is unclear. In young mice, MΦ depletion impairs muscle regeneration (Summan et al., 2006) and augmenting the MΦ response enhances myofiber regeneration (Rybalko, Hsieh, Merscham-Banda, Suggs, & Farrar, 2015). Therefore, greater MΦ infiltration of aged muscle following sterile injury could be a compensatory response to overcome local defects in MuSC function or a natural protective response to collateral damage of healthy tissue inflicted by the greater initial PMN influx. Further studies are needed to better clarify the role of infiltrating MΦ populations within the context of impaired regenerative capacity of aging muscle.

We found that aged mice displayed marked defects in muscle regeneration, but that RvD1 treatment had minimal impact on this response. This is in contrast to the beneficial effects of RvD1 on regenerating muscle fiber size seen in young healthy mice with this same dosing regimen (Markworth et al., 2020). Recent reports by others also show that local intramuscular injection of a distinct but related SPM, resolvin D2 (RvD2), can stimulate skeletal muscle regeneration in MuSC deficient (irradiated) mice (Giannakis et al., 2019). RvD1 treatment did reduce intramuscular MuSC number at day 5 post-injury in the current study, which is consistent with the effect observed previously in young mice (Markworth et al., 2020). However, in aged muscle this reduction in MuSC number was not accompanied by a parallel increase in the size of newly formed myofibers that was seen in young mice in our prior study. We previously interpreted this reduction in MuSCs to RvD1 driving differentiation and/or fusion of these myogenic precursors with regenerating myofibers (Markworth et al., 2020). However, MuSCs are well established to be indispensable for myofiber regeneration and aged mice already mounted a lower MuSC response when compared to young mice in the absence of RvD1 treatment. Therefore, the further reduction in MuSC number in aged mice with RvD1 treatment could also be interpreted as a deleterious effect of SPMs on MuSC proliferation, as is seen with systemic NSAID treatment (Mikkelsen et al., 2009). Unlike many prior studies reporting a deleterious effect of NSAIDs of myofiber regeneration following injury in mice however (Almekinders & Gilbert, 1986; Bondesen, Mills, Kegley, & Pavlath, 2004; Bryant et al., 2017; Dearth et al., 2016; Mishra, Friden, Schmitz, & Lieber, 1995; Shen, Li, Tang, Cummins, & Huard, 2005), we also saw no evidence of any negative effects of RvD1 on the formation or growth of regenerating myofibers within aged muscle in the current study, which would argue against this interpretation. Collectively these data suggest that RvD1 treatment may indeed have accelerated myogenic commitment, differentiation, and/or fusion of proliferated MuSCs during muscle regeneration in aged mice also, but that inherent defects within aged myofibers may prevent this from translating to a clear hypertrophy of regenerating myofibers that was seen in young mice (Markworth et al., 2020).

Aging is well-established to limit recovery of muscle function following injury (Brooks & Faulkner, 1990). Indeed, we found that aged mice showed marked deficits in recovery of absolute muscle strength (P_o_). This was partially due to an impaired ability of aged mice to recover their pre-injury muscle size, but also reflected a substantial deficit in relative strength (sP_o_) when normalized by their smaller muscle cross-sectional area. As a consequence, TA muscles from aged mice were not only of poorer quality than young muscles prior to injury, but also displayed maladaptive tissue remodeling that further worsened their relative contractile function during regeneration. Although treatment of aged mice with RvD1 did not impact upon recovery of P_o_, it did improve recovery of sP_o_. We could not attribute this to any effect of RvD1 on any cellular indices of myofiber regeneration, but RvD1 treatment did reduce the accumulation of non-myofiber (e.g. fibrotic and/or fatty) tissue within regenerating muscles of aged mice. Collectively, these data show that while systemic RvD1 treatment appears unable to overcome age-related defects in myofiber regeneration, it nonetheless limited maladaptive tissue remodeling and thus improved the quality of the regenerated muscle resulting in improved contractile function.

In conclusion, aging is associated with a local deficiency of intramuscular anti-inflammatory and pro-resolving lipid mediators which is related to both chronic low-grade muscle inflammation and heightened acute myeloid cell responses to muscle injury. Short term systemic administration of RvD1 as an immunoresolvent can limit excessive expression of inflammatory mediators, modulate the MuSC response, and limit maladaptive muscle tissue remodeling following muscle injury, but appears unable to overcome the marked age-associated defects in myofiber regenerative capacity in elderly mice. Given their key role in tissue regeneration (Serhan et al., 2012), it remains possible that other distinct families of SPMs such as the maresins which were deficient in aged muscle following injury could also contribute to defective myofiber regeneration in aging.

## Experimental Procedures

### Animals

Aged female C57BL/6 mice were obtained from the National Institute of Aging (NIA) at 20-22 mo, housed at the University of Michigan animal facilities for ~6 months, and utilized for experiments between 26-28 mo. Adult (4-6 mo) female C57BL/6 mice were obtained from Charles River Laboratories and served as young controls. All mice were housed under specific pathogen free conditions with ad-libitum access to food and water.

### Muscle Injury

Mice were anesthetized with 2% isoflurane and received bilateral intramuscular injection of the tibialis anterior (TA) muscle with 50 μL per limb of 1.2% barium chloride (BaCl_2_) in sterile saline to induce myofiber injury. Mice were returned to their home cage to recover and monitored until ambulatory with free access to food and water.

### Immunoresolvent Treatment

Resolvin D1 (RvD1) was purchased from Cayman Chemicals (10012554). Single use aliquots of RvD1 were prepared in amber glass vials (ThermoFisher, C4010-88AW) which were purged with nitrogen gas and stored at −80°C. On the day of use, the ethanol solution was evaporated to dryness under a gentle stream of nitrogen gas and RvD1 was re-suspended in sterile saline containing 0.1% ethanol. RvD1 stocks were handled in a darkroom and aqueous solutions of RvD1 were protected from light and used within 30 minutes of preparation. Aged mice were randomized to receive daily 100 μL intraperitoneal (IP) injections of either 100 ng of RvD1 or vehicle control (0.1% ethanol). The first dose was given ~5 min prior to muscle injury. Mice were allowed to recover for up to two-weeks with daily IP injection of 100 ng of RvD1 or vehicle control.

### Muscle Tissue Collection

Animals were euthanized via induction of bilateral pneumothorax while under deep isoflurane anesthesia. Muscles were rapidly dissected, blotted dry, weighed, and snap frozen in liquid nitrogen for molecular analysis. Muscles for histological analysis were cut transversely at the mid-belly, oriented longitudinally on a plastic support, covered with a thin layer of optimal cutting temperature (OCT) compound, and rapidly frozen in isopentane cooled on liquid nitrogen. Samples were stored at −80°C until analysis.

### Histology

Cross-sections (10 μm) were cut from the muscle mid-belly in a cryostat at −20°C and adhered to SuperFrost Plus slides. Tissue sections were air dried and then either stained with hematoxylin and eosin (H & E) or prepared for immunohistochemistry analysis. Slides for muscle immune cell staining were fixed in ice-cold acetone at −20°C and then air dried. Satellite cell staining slides were fixed in 4% paraformaldehyde (PFA), quenched with hydrogen peroxide, and antigen retrieval performed. Unfixed tissue sections were used for muscle fiber type staining. Prepared slides were blocked in 10% normal goat serum (GS, Invitrogen 10000C) or Mouse on Mouse (M.O.M) blocking reagent (Vector Laboratories, MKB-2213) as appropriate prior to overnight incubation at 4°C with primary antibodies including embryonic myosin heavy chain (DSHB, F1.652s, 1:20), Pax7 (DSHB, Pax7c, 1:100), myosin heavy chain (MHC) type I (DSHB, BA-D5c, 1:100), MHC IIa (DSHB, SC-71c, 1:100), MHC IIb (DSHB, BF-F3c, 1:100), laminin (Abcam, ab7463, 1:200), rat anti-mouse Ly-6G (Gr-1) (BD Biosciences, BD550291, 1:50), rat anti-mouse CD68 (Bio-Rad, MCA1957, 1:50), and rabbit polyclonal CD163 (Santa Cruz, sc-33560, 1:50). The following day, slides were incubated with either Alexa Fluor conjugated secondary antibodies (Invitrogen, 1:500 in PBS) or a Tyramide SuperBoost Kit as per the manufacturers instructions (Invitrogen, B40913) (for Pax7 only). Fluorescent dyes including 4′,6-diamidino-2-phenylindole (DAPI, Invitrogen, D21490, 2 μg/mL), wheat germ agglutinin (WGA) Alexa Fluor 647 conjugate (Invitrogen, W32466, 5 μg/mL), WGA CF405S conjugate (Biotium, 29027, 100 μg/mL), and phalloidin (Invitrogen, ActinRed 555 ReadyProbes, R37112) were used to counterstain cell nuclei, extracellular matrix, and muscle fibers respectively in particular experiments. Following washing in PBS slides were mounted using Fluorescence Mounting Medium (Agilent Dako, S302380) and fluorescent images were captured using a Nikon A1 confocal microscope.

### Image Analysis

Muscle tissue morphology was analyzed on stitched panoramic images of the entire muscle cross-section by high-throughput fully automated image analysis with the MuscleJ plugin for FIJI/Image J (Mayeuf-Louchart et al., 2018). Immune cells and satellite cells were manually counted throughout the entire muscle cross-section and then normalized to total tissue area as determined by MuscleJ. In all cases, the experimenter was blinded to the experimental group.

### Muscle Force Testing

Mice were anesthetized with 2% isoflurane and placed on a heated platform. The distal half of the TA muscle was isolated by dissecting the overlying skin and fascia. The knee joint was immobilized and a 4–0 silk suture tied around the distal TA tendon which was severed from its boney insertion and tied to the lever arm of a servomotor (6650LR, Cambridge Technology). A saline drip warmed to 37°C was continuously applied to the exposed muscle. The peroneal nerve was stimulated with 0.2 ms pulses using platinum electrodes with the stimulation voltage and muscle length adjusted to obtain optimal muscle length (L_o_) and maximum isometric twitch force (P_t_). The TA was then stimulated at increasing frequencies while held at L_o_ until maximum isometric tetanic force (P_o_) was achieved. One-minute rest was allowed between each tetanic contraction. Muscle length was then measured with calipers and optimum fiber length (L_f_) determined by multiplying L_o_ by the TA muscle L_f_/L_o_ ratio of 0.6 (Burkholder, Fingado, Baron, & Lieber, 1994). The cross-sectional area (CSA) of the muscle was calculated by dividing muscle mass by the product of L_f_ and 1.06 mg/mm^3^ (Mendez and Keys, 1960). Specific P_o_ (sP_o_) was calculated by dividing P_o_ by muscle CSA.

### RNA extraction and cDNA synthesis

Muscle samples were homogenized in TRIzol reagent and total RNA was isolated by Phenol/Chloroform extraction. RNA yield was determined using a NanoDrop Spectrophotometer (Nanodrop 2000c). Genomic DNA was removed by incubation with DNase I (Ambion, AM2222) followed by its heat inactivation. Total RNA (1 μg) was reversed transcribed to cDNA using SuperScript™ VILO™ Master Mix (Invitrogen, 11-755-050).

### Real time RT-qPCR

Whole muscle bulk gene expression was measured by RT-qPCR on a CFX96 Real-Time PCR Detection System (Bio-Rad, 1855195) in 20 μL reactions of iTaq™ Universal SYBR^®^ Green Supermix (Bio-Rad, #1725124) with 1 μM forward and reverse primers (Table 1). Relative mRNA expression was determined using the 2^−ΔΔ^Ct method. Primers are listed in Table 1.

**Table 1:**
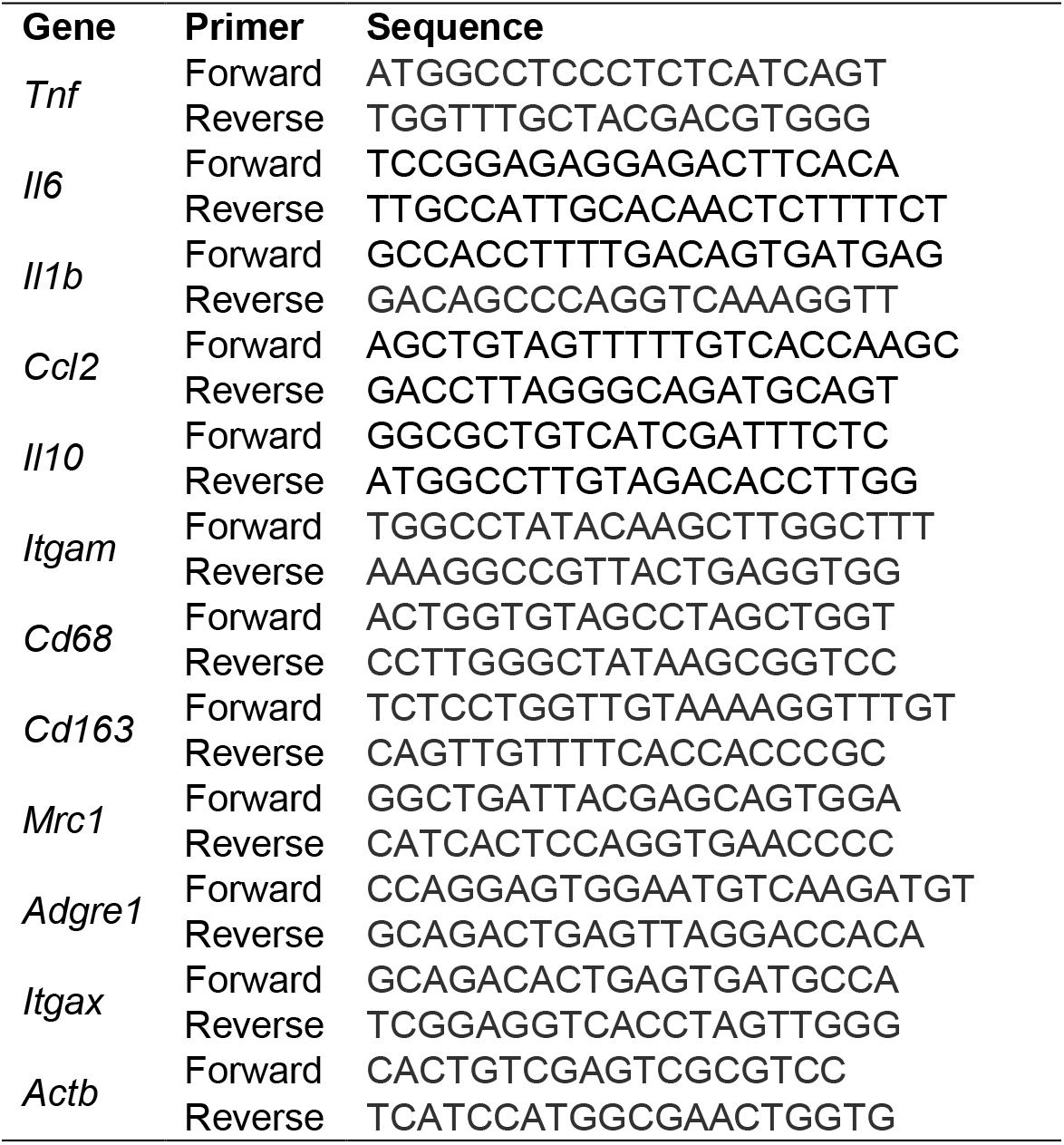
Real-time PCR primers

### Flow Cytometry Analysis of Basal Muscle Inflammation

The entire hind-limb musculature from each mouse was dissected, pooled, finely minced, and digested for 30 min at 37°C in Dulbecco’s Modified Eagle’s Medium (DMEM, Gibco, 11995-073) supplemented with 10 mg/mL collagenase II and 20 mg/mL dispase. The resulting digest solution was deactivated by adding fetal bovine serum (FBS) and then filtered through a 70-μm cell strainer and centrifuged at 450 × g for 5 min. Cells were Fc blocked with a CD16/CD32 antibody (ThermoFisher, 14-0161-82) for 10 minutes at 4°C and then incubated for 30 min at room temperature with primary antibodies including CD45-BV785 (BioLegend, 103149) CD11b-PE (BioLegend, 12-0112-82), CD68-FITC (BioLegend 137006), CD206-PE/Dazzle (BioLegend, 141732), and MHCII-APC/Cy7 (BioLegend, 107628). Following primary antibody incubation, cells were re-suspended in propidium iodide (PI, 2μL/mL) as a viability dye for 1 minute on ice (Life Technologies P3566). PI was then quenched with twice the volume of staining buffer and centrifuged at 450 × g for 1 minute. Flow cytometry analysis was performed using a Bio-Rad Ze5 Flow Cytometer and data was analyzed with FlowJo 10 software.

### Flow Cytometry Analysis of Acute Muscle Inflammation

At day 3 post- muscle injury both TA muscles from each mouse were pooled, finely minced, and digested for 60 min at 37°C for in Hank’s balanced salt solution lacking calcium and magnesium (HBSS^−/−^, Fisher Scientific), supplemented with 250 U/mL collagenase II, 4.0 U/mL dispase, and 2.5 mmol/L CaCl_2_. The resulting digest solution was filtered through a 40-μm cell strainer and centrifuged at 350 × g for 5 min. Cells were Fc blocked with a CD16/CD32 antibody (ThermoFisher, 14-0161-82) for 10 minutes at 4°C and then incubated for 30 min on ice with primary antibodies including CD45-PE (30-F11) (Thermo Fisher Scientific, 12-0451-82), Ly-6G-FITC (1A8) (BD Pharmingen, 551460), CD64-APC (X54-5/7.1) (BioLegend, 139306), CD11c-APCe780 (N418) (BioLegend, 47-0114-80), and Live/Dead fixable violet (Thermo Fisher Scientific, L34963). Flow cytometry was performed using a LSRFortessa cell analyzer (BD Biosciences) and data were analyzed with FlowJo 10 software.

### Metabolipidomic Profiling of Muscle Tissue

Muscle homogenates were analyzed by LC-MS/MS based lipidomic profiling following SPE extraction as previously described (Maddipati et al., 2014; Maddipati & Zhou, 2011; Markworth et al., 2020; Markworth et al., 2016; Markworth et al., 2013; Vella et al., 2019). Muscle samples were mechanically homogenized in 1 mL of 50 mM phosphate, pH 7.4 with 0.9% saline (PBS) using a bead mill with reinforced tubes and zirconium beads (Precellys). The tissue homogenates were centrifuged at 3,000 × g for 5 min and the supernatant was collected. Supernatants (0.85 ml) were spiked with 5 ng each of 15(S)-HETE-d8, 14(15)-EpETrE-d8, Resolvin D2-d5, Leukotriene B4-d4, and Prostaglandin E1-d4 as internal standards (in 150 μl methanol) for recovery and quantitation and mixed thoroughly. The internal standard spiked samples were applied to conditioned C18 cartridges, washed with 15% methanol in water followed by hexane and then dried under vacuum. The cartridges were eluted with 2 × 0.5 ml methanol with 0.1% formic acid. The eluate was dried under a gentle stream of nitrogen. The residue was re-dissolved in 50 μl methanol-25 mM aqueous ammonium acetate (1:1) and subjected to LC-MS/MS analysis.

HPLC was performed on a Prominence XR system (Shimadzu) using Luna C18 (3 μm, 2.1 × 150 mm) column. The mobile phase consisted of a gradient between A: methanol-water-acetonitrile (10:85:5 v/v) and B: methanol-water-acetonitrile (90:5:5 v/v), both containing 0.1% ammonium acetate. The gradient program with respect to the composition of B was as follows: 0-1 min, 50%; 1-8 min, 50-80%; 8-15 min, 80-95%; and 15-17 min, 95%. The flow rate was 0.2 ml/min. The HPLC eluate was directly introduced to the electrospray ion source of QTRAP5500 mass analyzer (ABSCIEX) in the negative ion mode with following conditions: Curtain gas: 35 psi, GS1: 35 psi, GS2: 65 psi, Temperature: 600 °C, Ion Spray Voltage: −1500 V, Collision gas: low, Declustering Potential: −60 V, and Entrance Potential: −7 V. The eluate was monitored by Multiple Reaction Monitoring (MRM) method to detect unique molecular ion – daughter ion combinations for each of the lipid mediators using a scheduled MRM around the expected retention time for each compound. Optimized Collisional Energies (18 – 35 eV) and Collision Cell Exit Potentials (7 – 10 V) were used for each MRM transition. Spectra of each peak detected in the scheduled MRM were recorded using Enhanced Product Ion scan to confirm the structural identity. The data were collected using Analyst 1.7 software and the MRM transition chromatograms were quantitated by MultiQuant software (both from ABSCIEX). The internal standard signals in each chromatogram were used for normalization, recovery, as well as relative quantitation of each analyte.

LC-MS data was analyzed using MetaboAnalyst 4.0 (Chong et al., 2018). Analytes with >50% missing values were removed from the data set and remaining missing values were replaced with half of the minimum positive value in the original data set. Heat maps were generated using the Pearson distance measure and the Ward clustering algorithm following autoscaling of features (analytes) without data transformation. Unsupervised principle component analysis (PCA) and volcano plots were generated in R using the FactoMineR and EnhancedVolcano packages respectively. Targeted statistical analysis was performed on pooled analyte concentrations from major lipid mediator biosynthetic pathways.

### Bone Marrow Derived Macrophage Culture

To confirm the established bioactivity of RvD1 on myeloid cells we performed assays of macrophage phagocytosis. Bone marrow was collected from tibias and femurs of young (4-6 mo) or aged (26-28 mo) female C57BL/6 mice and cultured for 7 days at 37°C and 5% CO_2_ in DMEM (Gibco, 11995-073) supplemented with 10% FBS, antibiotics, and 20 ng/mL recombinant murine GM-CSF (Biolegend, 576304). Cells were washed with PBS to remove non-adherent cells and adherent BMMs detached by incubation in TrypLE Select at 37°C (Gibco, 12563011) followed by gentle cell scraping. BMMs were plated into 96 well plates at a density of 1 × 10^5^ cells/well in growth media lacking GM-CSF and allowed to adhere overnight before use. MΦ were pre-treated for 15 min with RvD1 (1-100 nM) or vehicle control (0.1% ethanol) in growth media which was then replaced with pHrodo Green *E. Coli* Bio Particles (Invitrogen, P35366) prepared in hanks balanced salt solution containing calcium and magnesium (HBSS^+/+^) and incubated at 37°C for 1 hour in the continued presence of RvD1. Non-engulfed *E. coli* BioParticles were removed by washing with HBSS^+/+^ and intracellular fluorescence measured for the entire culture well using a plate reader at an excitation/emission of 509/533 nm.

### Statistics

Data is presented as the mean ± SEM. Statistical analysis was performed in GraphPad Prism 7. Between group differences were tested by two-tailed unpaired students t-tests (2 groups) or by a one-way analysis of variance (ANOVA) followed by either pair-wise Least Significance Difference (LSD) (3 groups) or Holm-Sidak post-hoc tests (>3 groups). p≤0.05 was used to determine statistical significance.

### Study approval

All animal experiments were approved by the University of Michigan Institutional Animal Care and Use committee (IACUC) (PRO00008744).

## Acknowledgments

This work was supported by the Glenn Foundation for Medical Research Post-Doctoral Fellowship in Aging Research (JFM), the National Institutes of Health (NIH) under the awards R01 (AG050676) (SVB), PO1 (AG051442) (SVB), P30 (AR069620) (CAA, SVB) and S10 (RR027926) (KRM), together with the 3M Foundation (CAA), American Federation for Aging Research (CAA), the University of Michigan Geriatrics Center/National Institute of Aging P30 (AG024824) (CAA, SVB), the University of Michigan Biomedical Engineering Department (CAA), and the Department of Defense and Congressionally Directed Medical Research Program W81XWH-20-1-0336 (CAA). The authors acknowledge members of the Benjamin Levi Laboratory (at the University of Michigan) for helpful scientific discussions.

## Author contributions

J.F.M and S.V.B conceived the study. S.V.B, K.R.M, P.C.D.M and C.A.A supervised the work. J.F.M, L.A.B, C.A.A, and S.V.B designed the experiments. J.F.M, L.A.B, E.L, J.L, J.A.C-M, and C.D performed the experiments. J.F.M, L.A.B, J.L, and K.R.M analyzed the data. J.F.M prepared the figures and wrote the manuscript with input from all authors.

## Data Availability Statement

Original data will be made available upon request.

---

**Supplemental Figure 1.**
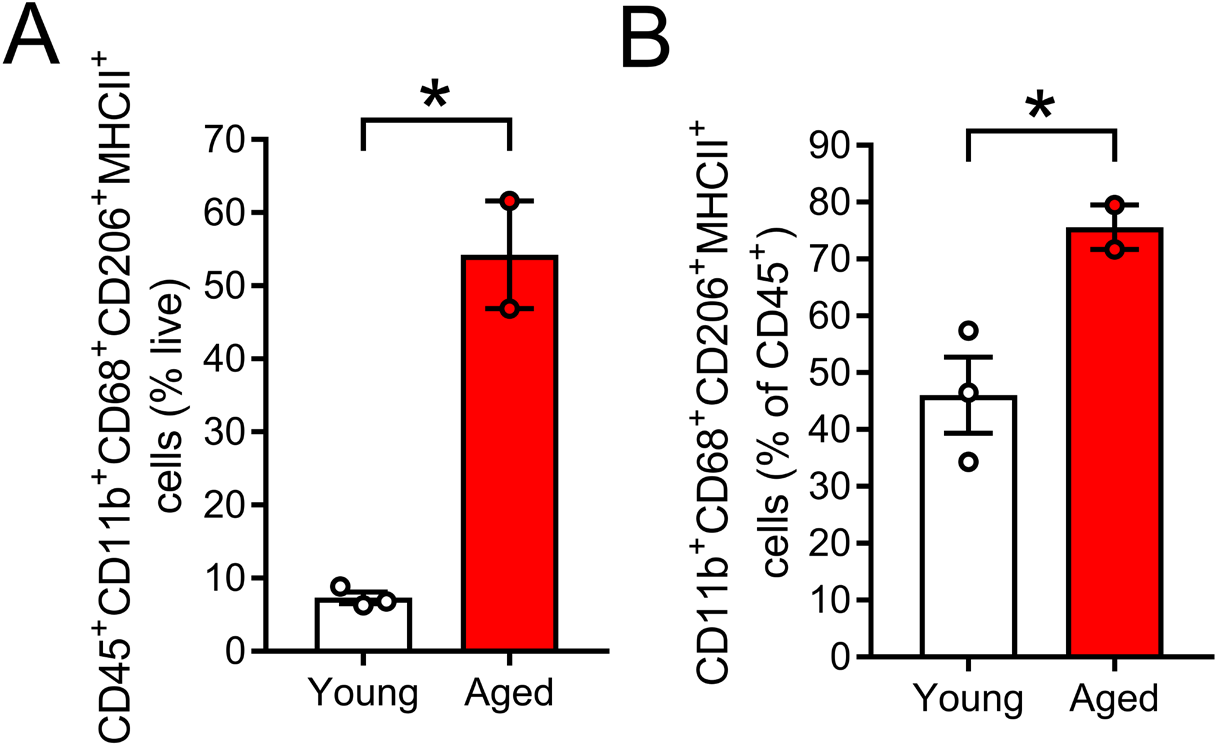
Flow cytometry analysis of basal myeloid cell accumulation in aged skeletal muscle: Single-cells isolated from pooled hind-limb muscles obtained from young (4-6 mo) and aged 26-28 mo) female C57BL/6 mice were analyzed by flow cytometry using antibodies including CD45-BV785, CD11b-PE, CD68-FITC, CD206-PE/Dazzle 594, MHCII (I-A/I-E)-APC/Cyanine 7. Dead cells were excluded from analysis with Propidium iodide (PI) staining. A: Quantification of M2-like resident muscle macrophages (defined as CD45^+^CD11b^+^CD68^+^CD206^+^MHCII^+^ cells) expressed as percentage of the total live cell population. B: Quantification of M2-like resident muscle macrophages (CD11b^+^CD68^+^CD206^+^MHCII^+^ cells) expressed as percentage of total intramuscular leukocytes (CD45^+^ cells). Bars show mean ± SEM of 2-3 mice per group with dots representing data from each individual mouse. *Denotes p<0.05 vs vehicle group by a two-tailed unpaired t-test.

**Supplemental Figure 2.**
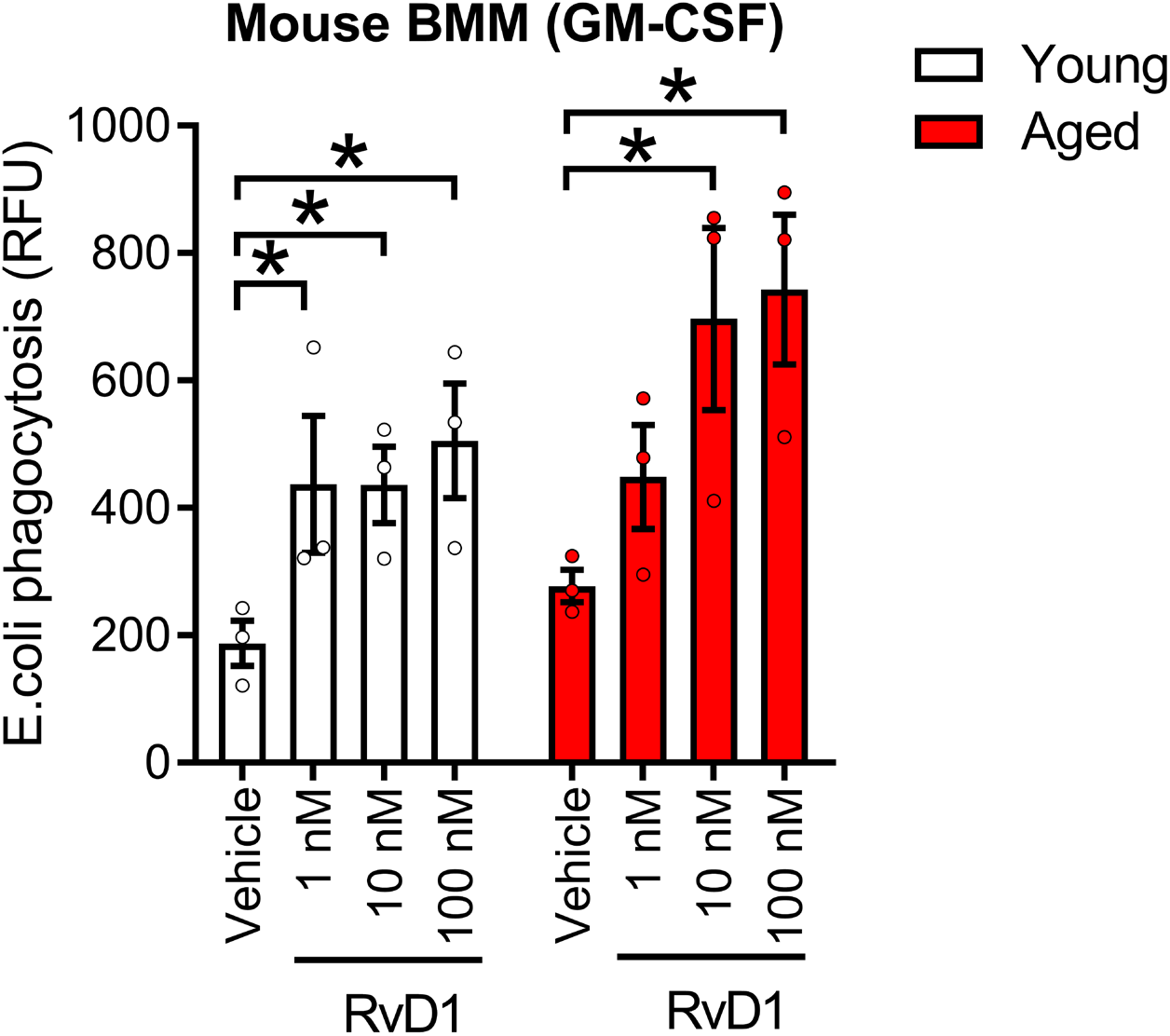
Resolvin D1 stimulates phagocytosis in macrophages from both young and aged hosts: A: Bone marrow-derived macrophages (BMM) from young (4-6 mo) and aged (26-28 mo) female C57BL/6 mice were obtained by culturing myeloid precursor cells for 7 days in the presence of 20 ng/mL GM-CSF. Young and aged GM-CSF derived BMMs were then pre-treated for 15 minutes with resolvin D1 (RvD1, 1-100 nM) prior to incubation with pHrodo Green E. Coli Bio Particles for 60 min at 37°C. Phagocytosis was quantified as the fluorescence intensity as determined by measurement of the entire well with a fluorescent plate reader. Bars show the mean ± SEM of cells from three young and aged donor mice for each experimental condition. *Denotes p<0.05 by two-way ANOVA with pair-wise Holm-Sidak post-hoc tests.

**Supplemental Figure 3.**
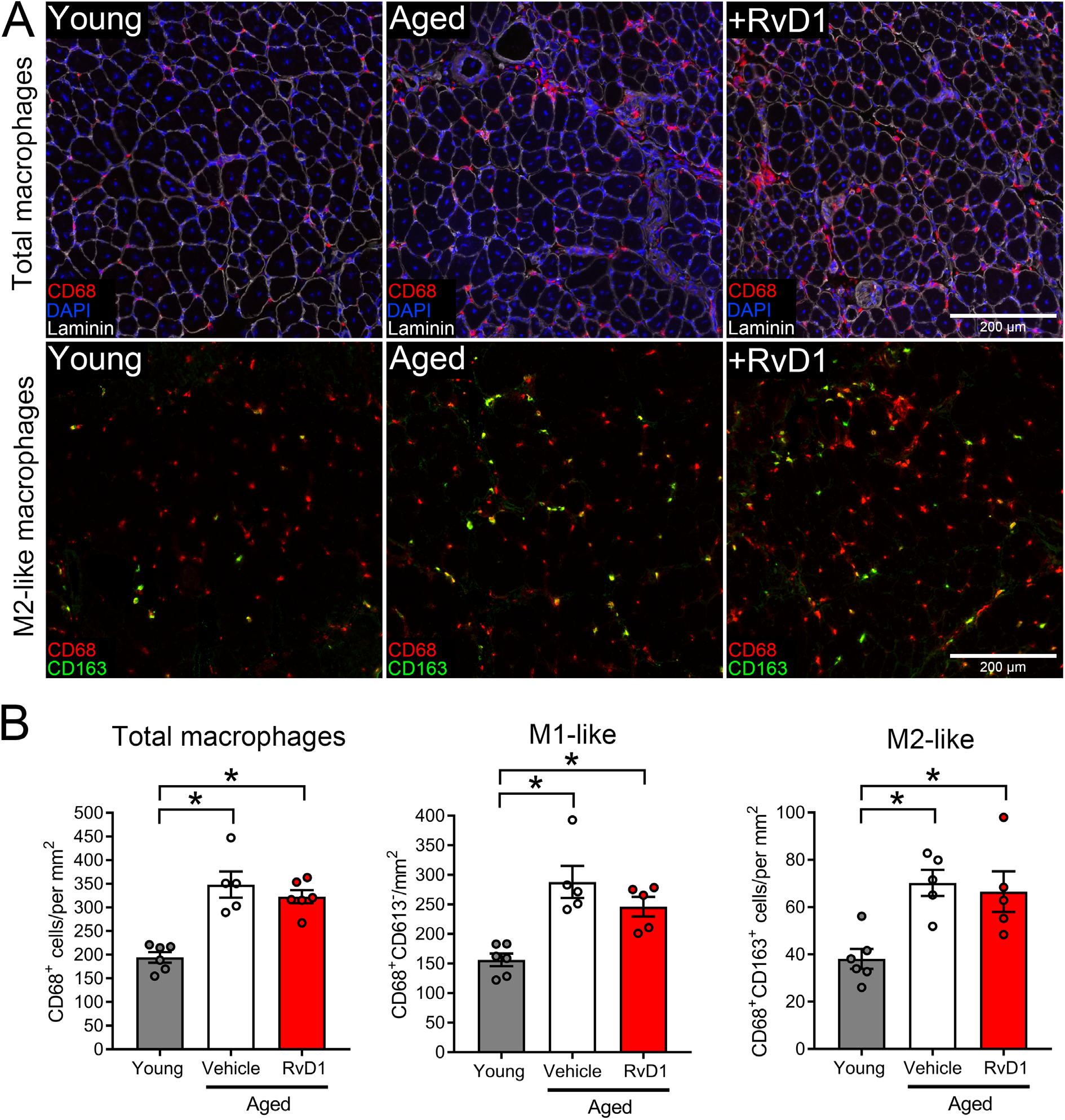
Resolvin D1 does not impact persistent elevation of intramuscular macrophages following muscle injury in aged mice. A: Young (4-6 mo) and aged (26-28 mo) female C57BL/6 mice received bilateral intramuscular injection of the tibialis anterior (TA) muscle with 50 μL of 1.2% barium chloride (BaCl_2_) to induce myofiber injury. Aged mice were then treated with daily intraperitoneal (IP) injections of resolvin D1 (RvD1) (100 ng) or vehicle (0.1% ethanol) for 14 days with the first dose administered ~5 min prior to injury. TA muscles were collected two-weeks later (day 14 post-injury) and tissue cross-sections were stained for total macrophages (MΦ, CD68), or M2-like MΦ (CD163). Cell nuclei and the basal lamina were counterstained with DAPI and a laminin antibody respectively. Scale bars are 200 μm. B: Quantification of myeloid cells in regenerating muscle including total MΦ (CD68^+^ cells), M1-like MΦ (CD68^+^CD163^−^ cells), and M2-like MΦ (CD68^+^CD163^+^ cells). Bars show mean ± SEM of 5-6 mice per group with dots representing data from each individual mouse. Denotes *p<0.05 by one-way ANOVA with pairwise LSD post-hoc tests.

**Supplemental Table 1A:** Cyclooxygenase (COX) metabolite concentration (pg/mg) in young and aged mouse tibialis anterior (TA) muscles

| Pathway | Substrate | Analyte | Uninjured |  | Day 1 post-injury |  | Day 3 post-injury |  | Day 5 post-injury |  |
| --- | --- | --- | --- | --- | --- | --- | --- | --- | --- | --- |
|  |  |  | Young | Aged | Young | Aged | Young | Aged | Young | Aged |
| Cyclooxygenase | 20:3n-6 | PGE1 | ND ± ND | 0.05 ± 0.04 | 0.29 ± 0.13 | 0.15 ± 0.08 | 4.91 ± 0.69 | 1.18 ± 0.50 | 2.98 ± 0.38 | 0.81 ± 0.39 |
|  | 20:3n-6 | PGF1a | 0.47 ± 0.05 | 0.35 ± 0.02 | ND ± ND | 0.20 ± 0.07 | 0.65 ± 0.13 | 0.55 ± 0.04 | 0.53 ± 0.14 | 0.66 ± 0.08 |
|  | 20:3n-6 | 15-keto PGE1 | 0.02 ± 0.01 | ND ± ND | ND ± ND | ND ± ND | ND ± ND | ND ± ND | ND ± ND | ND ± ND |
|  | 20:3n-6 | 13,14dhPGE1 | 0.09 ± 0.08 | 0.07 ± 0.06 | ND ± ND | ND ± ND | ND ± ND | 0.05 ± 0.03 | 0.13 ± 0.12 | ND ± ND |
|  | 20:3n-6 | 13,14dhPGE1 | ND ± ND | 0.18 ± 0.02 | ND ± ND | 0.06 ± 0.03 | 0.23 ± 0.06 | 0.29 ± 0.06 | 0.18 ± 0.08 | 0.29 ± 0.07 |
|  | 20:3n-6 | D17-PGE1 | ND ± ND | ND ± ND | ND ± ND | ND ± ND | ND ± ND | ND ± ND | ND ± ND | ND ± ND |
|  | 20:3n-6 | Bicyclo PGE1 | ND ± ND | ND ± ND | 0.17 ± 0.11 | 0.08 ± 0.07 | 0.13 ± 0.06 | 0.23 ± 0.06 | ND ± ND | ND ± ND |
|  | 20:3n-6 | 6-keto PGE1 | ND ± ND | ND ± ND | ND ± ND | ND ± ND | ND ± ND | ND ± ND | ND ± ND | ND ± ND |
|  | 20:3n-6 | 2,3-dinor PGE1 | ND ± ND | ND ± ND | ND ± ND | 0.06 ± 0.05 | ND ± ND | ND ± ND | ND ± ND | ND ± ND |
|  | 20:3n-6 | 19(R)-hydroxy PGE1 | ND ± ND | ND ± ND | ND ± ND | ND ± ND | ND ± ND | ND ± ND | ND ± ND | ND ± ND |
|  | 20:3n-6 | 15(R)-PGE1 | ND ± ND | ND ± ND | 0.06 ± 0.05 | ND ± ND | ND ± ND | 0.40 ± 0.19 | 0.11 ± 0.09 | 0.21 ± 0.12 |
|  | 20:4n-6 | TXB2 | 8.81 ± 1.52 | 7.36 ± 1.56 | 4.15 ± 0.78 | 4.07 ± 0.31 | 17.19 ± 2.89 | 16.66 ± 2.68 | 12.51 ± 1.94 | 14.15 ± 1.64 |
|  | 20:4n-6 | 12(S)-HHTrE | 13.75 ± 0.86 | 11.29 ± 1.60 | 8.31 ± 0.93 | 11.56 ± 0.98 | 36.47 ± 5.81 | 41.82 ± 8.11 | 42.63 ± 3.52 | 47.67 ± 8.63 |
|  | 20:4n-6 | PGD2 | 5.40 ± 0.26 | 6.28 ± 1.14 | 4.00 ± 0.35 | 4.49 ± 0.56 | 119.32 ± 26.33 | 60.80 ± 14.61 | 79.03 ± 10.63 | 39.07 ± 13.83 |
|  | 20:4n-6 | PGE2 | 18.93 ± 1.43 | 21.86 ± 3.20 | 18.23 ± 1.92 | 19.03 ± 1.67 | 101.84 ± 12.58 | 76.99 ± 14.86 | 96.72 ± 15.01 | 82.08 ± 17.91 |
|  | 20:4n-6 | PGF2a | 12.51 ± 0.89 | 11.29 ± 1.52 | 4.00 ± 0.19 | 5.59 ± 0.76 | 13.82 ± 2.43 | 17.13 ± 2.32 | 15.33 ± 2.01 | 18.91 ± 2.98 |
|  | 20:4n-6 | 6kPGF1a | 2.72 ± 0.32 | 2.82 ± 0.46 | 1.87 ± 0.28 | 1.82 ± 0.16 | 11.00 ± 1.40 | 8.15 ± 0.97 | 32.99 ± 4.50 | 18.24 ± 2.71 |
|  | 20:4n-6 | PGJ2 | 0.23 ± 0.06 | 0.31 ± 0.07 | 0.38 ± 0.04 | 0.33 ± 0.07 | 9.31 ± 1.30 | 5.25 ± 1.25 | 10.19 ± 2.14 | 5.05 ± 1.74 |
|  | 20:4n-6 | PGA2 | 0.08 ± 0.07 | 0.33 ± 0.17 | 0.41 ± 0.14 | 0.51 ± 0.12 | 4.93 ± 2.58 | 4.28 ± 1.99 | 7.89 ± 3.03 | 4.54 ± 2.20 |
|  | 20:4n-6 | 15-keto PGE2 | 1.32 ± 0.18 | 1.25 ± 0.23 | 1.30 ± 0.19 | 1.17 ± 0.21 | 4.54 ± 0.76 | 2.24 ± 0.52 | 3.75 ± 0.68 | 2.43 ± 0.68 |
|  | 20:4n-6 | 15-keto PGF2a | 0.05 ± 0.04 | 0.14 ± 0.07 | 0.05 ± 0.04 | 0.05 ± 0.03 | 3.24 ± 0.67 | 1.62 ± 0.42 | 2.17 ± 0.32 | 1.19 ± 0.40 |
|  | 20:4n-6 | 13,14dh-15k-PGE2 | 1.42 ± 0.13 | 1.58 ± 0.26 | 0.65 ± 0.49 | 0.20 ± 0.19 | 4.37 ± 0.59 | 3.17 ± 0.57 | 2.34 ± 0.90 | 3.60 ± 0.71 |
|  | 20:4n-6 | 13,14dh-15k-PGD2 | 1.61 ± 0.25 | 1.55 ± 0.36 | 1.00 ± 0.21 | 0.66 ± 0.05 | 6.91 ± 1.30 | 2.66 ± 0.59 | 6.70 ± 1.15 | 2.54 ± 0.75 |
|  | 20:4n-6 | 19(R)-OH PGF2a & 20-OH PGF2a | 0.22 ± 0.03 | 0.22 ± 0.02 | 0.29 ± 0.04 | 0.17 ± 0.01 | 0.31 ± 0.01 | 0.17 ± 0.05 | 0.36 ± 0.08 | 0.22 ± 0.06 |
|  | 20:4n-6 | 8-isoPGF2a & 11bPGF2a | 0.39 ± 0.02 | 0.40 ± 0.09 | 0.11 ± 0.05 | 0.33 ± 0.02 | 0.66 ± 0.22 | 1.14 ± 0.18 | 0.90 ± 0.13 | 1.10 ± 0.26 |
|  | 20:4n-6 | 11dh-2,3-dinor TXB2 | 0.82 ± 0.13 | 0.46 ± 0.16 | 0.23 ± 0.15 | ND ± ND | 0.30 ± 0.18 | ND ± ND | 0.38 ± 0.24 | ND ± ND |
|  | 20:4n-6 | D12-PGJ2 | 0.36 ± 0.04 | 0.38 ± 0.09 | 0.21 ± 0.05 | 0.21 ± 0.09 | 3.99 ± 2.31 | 2.68 ± 0.86 | 3.26 ± 1.05 | 1.26 ± 0.82 |
|  | 20:4n-6 | tetranor PGEM | ND ± ND | 0.01 ± 0.00 | ND ± ND | ND ± ND | ND ± ND | ND ± ND | ND ± ND | ND ± ND |
|  | 20:4n-6 | 15d-D12,14-PGJ2 | ND ± ND | ND ± ND | ND ± ND | ND ± ND | 0.15 ± 0.08 | 0.05 ± 0.03 | 0.06 ± 0.04 | ND ± ND |
|  | 20:4n-6 | Bicyclo PGE2 | ND ± ND | ND ± ND | 0.03 ± 0.02 | ND ± ND | 0.48 ± 0.05 | 0.22 ± 0.08 | 0.29 ± 0.13 | 0.19 ± 0.12 |
|  | 20:4n-6 | 19(R)-OH PGE2 & 20-OH PGE2 | ND ± ND | ND ± ND | ND ± ND | ND ± ND | ND ± ND | ND ± ND | ND ± ND | ND ± ND |
|  | 20:4n-6 | 2,3-dinor TXB2 | ND ± ND | ND ± ND | ND ± ND | ND ± ND | ND ± ND | ND ± ND | ND ± ND | ND ± ND |
|  | 20:4n-6 | 11dh-TXB2 | ND ± ND | ND ± ND | 0.03 ± 0.02 | ND ± ND | 0.04 ± 0.02 | ND ± ND | 0.11 ± 0.06 | ND ± ND |
|  | 20:4n-6 | 13,14dh-15k-PGF2a | ND ± ND | ND ± ND | ND ± ND | ND ± ND | ND ± ND | ND ± ND | ND ± ND | 0.07 ± 0.05 |
|  | 20:4n-6 | 6,15-diketo PGFa | ND ± ND | ND ± ND | ND ± ND | ND ± ND | ND ± ND | ND ± ND | ND ± ND | ND ± ND |
|  | 20:4n-6 | iPF-VI | ND ± ND | ND ± ND | ND ± ND | 0.03 ± 0.02 | ND ± ND | 0.04 ± 0.02 | ND ± ND | ND ± ND |
|  | 20:5n-3 | TXB3 | 1.08 ± 0.26 | 0.68 ± 0.16 | 0.19 ± 0.08 | 0.07 ± 0.05 | 1.55 ± 0.44 | 0.95 ± 0.09 | 1.38 ± 0.32 | 0.46 ± 0.14 |
|  | 20:5n-3 | PGE3 | 0.23 ± 0.06 | 0.29 ± 0.09 | 0.14 ± 0.07 | 0.15 ± 0.07 | 2.38 ± 0.34 | 1.27 ± 0.24 | 1.90 ± 0.37 | 0.97 ± 0.33 |
|  | 20:5n-3 | PGF3a | 0.17 ± 0.01 | 0.20 ± 0.02 | 0.01 ± ND | ND ± ND | 0.11 ± 0.07 | 0.25 ± 0.02 | 0.16 ± 0.07 | 0.08 ± 0.04 |
|  | 20:5n-3 | PGD3 | ND ± ND | ND ± ND | ND ± ND | ND ± ND | 1.75 ± 0.49 | 0.50 ± 0.23 | 1.25 ± 0.31 | 0.32 ± 0.30 |
|  | 20:5n-3 | 11dh TXB3 | ND ± ND | ND ± ND | 0.01 ± ND | ND ± ND | ND ± ND | ND ± ND | ND ± ND | ND ± ND |
|  | 20:5n-3 | 15d-D12,14-PGJ3 | ND ± ND | ND ± ND | 0.01 ± ND | ND ± ND | ND ± ND | ND ± ND | ND ± ND | ND ± ND |
|  |  | Sum | 70.83 ± 3.97 | 69.56 ± 10.21 | 46.29 ± 4.50 | 51.22 ± 4.12 | 358.53 ± 50.23 | 250.98 ± 47.35 | 326.44 ± 43.45 | 246.36 ± 53.48 |

Values are mean ± SEM of 5-7 mice/group. ND = Below limits of detection of the assay.

**Supplemental Table 1B:** Lipoxygenase (LOX) metabolite concentration (pg/mg) in young and aged mouse tibialis anterior (TA) muscles

| Pathway | Substrate | Analyte | Uninjured |  | Day 1 post-injury |  | Day 3 post-injury |  | Day 5 post-injury |  |
| --- | --- | --- | --- | --- | --- | --- | --- | --- | --- | --- |
|  |  |  | Young | Aged | Young | Aged | Young | Aged | Young | Aged |
| 5-LOX | 20:4n-6 | 5-HETE | 9.27 ± 1.44 | 5.02 ± 0.68 | 16.91 ± 3.03 | 17.78 ± 4.25 | 48.31 ± 7.72 | 21.73 ± 5.54 | 37.03 ± 1.53 | 15.89 ± 1.78 |
|  | 20:4n-6 | 5-oxoETE | 3.03 ± 0.66 | 1.60 ± 0.22 | 4.72 ± 1.02 | 5.39 ± 1.31 | 15.36 ± 2.85 | 6.33 ± 1.68 | 10.25 ± 1.03 | 5.09 ± 0.71 |
|  | 20:4n-6 | 12-OxoLTB4 | 0.29 ± 0.10 | 0.08 ± 0.05 | 0.23 ± 0.07 | 0.23 ± 0.03 | 0.79 ± 0.25 | 0.49 ± 0.23 | 0.62 ± 0.26 | 0.23 ± 0.13 |
|  | 20:4n-6 | 5(S),15(S)-DihETE | ND ± ND | ND ± ND | ND ± ND | ND ± ND | ND ± ND | ND ± ND | ND ± ND | ND ± ND |
|  | 18:3n-3 | 5(S)-HETrE | 0.23 ± 0.01 | 0.11 ± 0.01 | 0.21 ± 0.03 | 0.23 ± 0.08 | 0.80 ± 0.13 | 0.40 ± 0.11 | 0.56 ± 0.04 | 0.27 ± 0.03 |
|  | 18:3n-3 | 9(S)-HOTrE | 4.73 ± 0.42 | 1.87 ± 0.23 | 1.85 ± 0.35 | 2.46 ± 0.69 | 3.23 ± 0.46 | 2.39 ± 0.50 | 4.56 ± 0.39 | 1.88 ± 0.21 |
|  | 18:3n-3 | 9-OxoOTrE | 2.50 ± 0.19 | 1.09 ± 0.19 | 0.81 ± 0.11 | 1.35 ± 0.48 | 1.83 ± 0.36 | 1.15 ± 0.23 | 2.92 ± 0.39 | 1.40 ± 0.17 |
|  | 22:6n-3 | 7-HDoHE | 1.57 ± 0.14 | 1.06 ± 0.15 | 7.22 ± 1.94 | 6.17 ± 2.63 | 11.86 ± 1.76 | 5.01 ± 0.98 | 5.24 ± 0.75 | 2.16 ± 0.41 |
|  | 20:5n-3 | 5-HEPE | 1.23 ± 0.10 | 1.02 ± 0.20 | 2.25 ± 0.34 | 2.12 ± 0.39 | 8.53 ± 1.37 | 3.77 ± 0.82 | 7.94 ± 0.41 | 2.29 ± 0.45 |
|  | 20:5n-3 | LTB5 | 0.45 ± 0.15 | 0.39 ± 0.10 | 0.08 ± 0.07 | ND ± ND | 0.40 ± 0.18 | ND ± ND | 0.24 ± 0.18 | ND ± ND |
| 5-LOX sum |  |  | 23.31 ± 2.16 | 12.07 ± 1.58 | 34.29 ± 6.10 | 35.78 ± 9.73 | 91.11 ± 14.41 | 41.35 ± 9.76 | 69.36 ± 3.21 | 29.28 ± 3.09 |
| 8-LOX | 20:5n-3 | 8-HEPE | 1.18 ± 0.09 | 0.64 ± 0.18 | 1.50 ± 0.24 | 1.19 ± 0.26 | 4.58 ± 0.68 | 2.55 ± 0.42 | 4.24 ± 0.34 | 1.41 ± 0.29 |
|  | 20:4n-6 | 8-HETE | 6.15 ± 0.37 | 2.43 ± 0.64 | 8.08 ± 1.24 | 9.40 ± 3.49 | 22.82 ± 3.97 | 12.98 ± 3.14 | 21.36 ± 1.88 | 10.53 ± 1.44 |
|  | 8-LOX sum |  | 7.33 ± 0.43 | 3.07 ± 0.76 | 9.58 ± 1.44 | 10.58 ± 3.74 | 27.40 ± 4.63 | 15.52 ± 3.51 | 25.59 ± 2.15 | 11.94 ± 1.68 |
| 12-LOX | 18:2n-6 | 9-HODE | 37.02 ± 2.58 | 25.93 ± 2.49 | 41.48 ± 2.80 | 70.72 ± 13.04 | 94.36 ± 10.78 | 72.14 ± 9.27 | 83.87 ± 4.67 | 90.90 ± 10.04 |
|  | 18:2n-6 | 9-OxoODE | 34.34 ± 4.14 | 29.46 ± 4.48 | 36.04 ± 7.12 | 50.09 ± 15.09 | 72.14 ± 10.05 | 42.38 ± 6.84 | 63.96 ± 10.16 | 56.76 ± 8.40 |
|  | 18:3n-3 | 8(S)-HETrE | 1.88 ± 0.23 | 0.83 ± 0.16 | 1.77 ± 0.31 | 1.50 ± 0.27 | 6.87 ± 1.08 | 2.28 ± 0.38 | 4.72 ± 0.47 | 1.93 ± 0.26 |
|  | 20:2n-6 | 11(R)-HEDE | 1.01 ± 0.05 | 0.82 ± 0.11 | 1.12 ± 0.16 | 1.42 ± 0.21 | 8.42 ± 0.83 | 4.23 ± 0.90 | 5.73 ± 0.42 | 2.90 ± 0.35 |
|  | 20:4n-6 | 12-HETE | 1256.59 ± 178.43 | 793.82 ± 157.77 | 888.38 ± 180.77 | 914.28 ± 80.08 | 2687.71 ± 475.07 | 2332.08 ± 452.58 | 2905.47 ± 384.39 | 2500.70 ± 243.84 |
|  | 20:4n-6 | 5(S),12(S)-DihETE | 26.26 ± 5.78 | 13.99 ± 3.26 | 11.97 ± 2.93 | 10.29 ± 2.11 | 34.71 ± 6.14 | 29.24 ± 3.82 | 30.16 ± 5.62 | 27.63 ± 7.03 |
|  | 20:4n-6 | tetranor 12-HETE | 2.00 ± 0.27 | 2.31 ± 0.41 | 0.64 ± 0.12 | 0.87 ± 0.30 | 1.60 ± 0.20 | 1.39 ± 0.25 | 2.78 ± 0.62 | 1.39 ± 0.19 |
|  | 20:4n-6 | 12-OxoETE | ND ± ND | ND ± ND | ND ± ND | ND ± ND | ND ± ND | ND ± ND | ND ± ND | ND ± ND |
|  | 20:5n-3 | 12-HEPE | 288.97 ± 58.63 | 200.85 ± 48.97 | 129.20 ± 27.68 | 112.41 ± 18.60 | 617.01 ± 100.74 | 465.44 ± 71.80 | 704.04 ± 96.70 | 361.08 ± 62.96 |
|  | 22:6n-3 | 14-HDoHE | 77.94 ± 19.48 | 51.49 ± 11.63 | 84.82 ± 16.17 | 58.66 ± 10.02 | 240.33 ± 39.99 | 107.64 ± 19.00 | 135.28 ± 11.81 | 79.92 ± 8.98 |
| 12-LOX sum |  |  | 1729.60 ± 259.02 | 1119.51 ± 213.80 | 1195.42 ± 225.26 | 1220.24 ± 109.20 | 3763.16 ± 625.65 | 3056.84 ± 551.45 | 3936.06 ± 483.74 | 3123.23 ± 286.91 |
| 15-LOX | 18:2n-6 | 13-HODE | 232.10 ± 17.49 | 150.40 ± 14.07 | 174.53 ± 15.62 | 265.96 ± 51.86 | 408.27 ± 40.98 | 307.41 ± 39.28 | 438.65 ± 15.39 | 341.74 ± 22.64 |
|  | 18:2n-6 | 13-OxoODE | 66.70 ± 4.50 | 43.83 ± 4.43 | 65.98 ± 8.88 | 112.80 ± 32.43 | 153.41 ± 21.80 | 90.07 ± 11.86 | 115.45 ± 13.97 | 88.99 ± 4.86 |
|  | 18:3n-3 | 13(S)-HOTrE | 5.15 ± 0.99 | 1.71 ± 0.41 | 2.00 ± 0.25 | 2.43 ± 0.53 | 3.33 ± 0.52 | 2.71 ± 0.39 | 7.25 ± 0.50 | 4.25 ± 0.58 |
|  | 18:3n-6 | 13(S)-HOTrE(g) | 0.47 ± 0.29 | 0.24 ± 0.15 | 0.37 ± 0.23 | 1.18 ± 0.54 | 0.86 ± 0.33 | 1.23 ± 0.50 | 1.71 ± 0.63 | 2.25 ± 0.65 |
|  | 20:2n-6 | 15(S)-HEDE | 0.35 ± 0.03 | 0.21 ± 0.02 | 0.31 ± 0.05 | 0.35 ± 0.09 | 2.00 ± 0.26 | 0.89 ± 0.18 | 1.35 ± 0.08 | 0.68 ± 0.06 |
|  | 20:2n-6 | 15-OxoEDE | ND ± ND | ND ± ND | 0.17 ± 0.07 | 0.25 ± 0.07 | 0.99 ± 0.21 | 0.47 ± 0.09 | 0.44 ± 0.12 | 0.13 ± 0.07 |
|  | 20:4n-6 | 15-HETE | 19.52 ± 1.65 | 11.90 ± 1.32 | 21.72 ± 2.98 | 24.82 ± 5.35 | 79.04 ± 10.53 | 48.93 ± 11.33 | 72.42 ± 3.23 | 48.31 ± 6.64 |
|  | 20:4n-6 | 15-OxoETE | 0.79 ± 0.11 | 1.26 ± 0.42 | 1.25 ± 0.21 | 1.39 ± 0.34 | 2.64 ± 0.67 | 1.61 ± 0.37 | 2.47 ± 0.34 | 1.48 ± 0.15 |
|  | 20:4n-6 | 8(S),15(S)-DihETE | 0.17 ± 0.05 | ND ± ND | ND ± ND | ND ± ND | ND ± ND | 0.15 ± 0.07 | 0.25 ± 0.09 | ND ± ND |
|  | 20:4n-6 | 5(S),15(S)-DihETE | ND ± ND | ND ± ND | ND ± ND | ND ± ND | ND ± ND | ND ± ND | ND ± ND | ND ± ND |
|  | 20:5n-3 | 15(S)-HEPE | 1.38 ± 0.20 | 0.68 ± 0.18 | 1.72 ± 0.27 | 1.33 ± 0.25 | 14.59 ± 3.36 | 6.74 ± 2.78 | 18.82 ± 2.10 | 3.61 ± 0.86 |
|  | 20:5n-3 | 5(S),15(S)-DIHEPE | 0.32 ± 0.10 | 0.12 ± 0.03 | ND ± ND | ND ± ND | ND ± ND | ND ± ND | 0.15 ± 0.07 | 0.17 ± 0.07 |
|  | 22:6n-3 | 17-HDoHE | 2.23 ± 0.62 | 2.10 ± 0.31 | 5.61 ± 1.37 | w ± 2.60 | 25.64 ± 4.45 | 10.59 ± 3.46 | 16.77 ± 1.97 | 6.90 ± 1.76 |
| 15-LOX sum |  |  | 324.55 ± 17.77 | 212.91 ± 19.43 | 273.72 ± 27.86 | 416.43 ± 92.53 | 690.82 ± 76.50 | 470.87 ± 64.86 | 653.39 ± 29.70 | 496.54 ± 34.46 |
| Total LOX sum |  |  | 2084.81 ± 273.61 | 1347.59 ± 227.59 | 1513.04 ± 248.67 | 1683.05 ± 184.01 | 4572.51 ± 697.20 | 3584.59 ± 621.34 | 4621.75 ± 461.56 | 3661.01 ± 318.71 |

Values are mean ± SEM of 5-7 mice/group. ND = Below limits of detection of the assay.

**Supplemental Table 1C:** Epoxygenase (CYP) metabolite concentration (pg/mg) in young and aged mouse tibialis anterior (TA) muscles

| Pathway | Substrate | Analyte | Uninjured |  | Day 1 post-injury |  | Day 3 post-injury |  | Day 5 post-injury |  |
| --- | --- | --- | --- | --- | --- | --- | --- | --- | --- | --- |
|  |  |  | Young | Aged | Young | Aged | Young | Aged | Young | Aged |
| Epoxygenase | 18:2n-6 | 9(10)-EpOME | 121.03 ± 6.84 | 19.24 ± 9.47 | 70.93 ± 10.40 | 54.08 ± 35.51 | 65.46 ± 23.19 | 41.71 ± 9.14 | 75.14 ± 15.18 | 28.94 ± 7.38 |
|  | 18:2n-6 | 12(13)-EpOME | 48.54 ± 4.10 | 20.31 ± 4.50 | 35.30 ± 4.83 | 44.19 ± 13.41 | 49.44 ± 7.20 | 25.12 ± 3.33 | 39.01 ± 4.14 | 17.97 ± 1.02 |
|  | 18:2n-6 | 9,10-DiHOME | 6.01 ± 0.54 | 4.85 ± 1.14 | 6.29 ± 0.51 | 6.32 ± 0.57 | 7.82 ± 0.76 | 7.97 ± 0.86 | 11.11 ± 0.79 | 8.60 ± 0.77 |
|  | 18:2n-6 | 12,13-DiHOME | 5.07 ± 0.37 | 4.82 ± 1.02 | 6.60 ± 0.69 | 6.35 ± 0.53 | 7.09 ± 0.74 | 7.92 ± 0.90 | 13.13 ± 1.45 | 9.95 ± 0.95 |
|  | 20:4n-6 | 5(6)-EpETRE | 9.26 ± 1.49 | 1.67 ± 0.50 | 5.56 ± 1.03 | 3.18 ± 0.87 | 5.66 ± 1.28 | 1.46 ± 0.35 | 2.18 ± 0.33 | 0.81 ± 0.08 |
|  | 20:4n-6 | 8(9)-EpETRE | 3.96 ± 0.80 | 1.11 ± 0.34 | 0.25 ± 0.24 | 0.38 ± 0.17 | 1.75 ± 0.63 | 2.77 ± 0.45 | 2.26 ± 0.52 | 2.29 ± 0.63 |
|  | 20:4n-6 | 11(12)-EpETRE | 9.89 ± 1.80 | 2.81 ± 0.76 | 10.52 ± 1.86 | 9.68 ± 2.86 | 22.37 ± 4.67 | 8.55 ± 2.07 | 13.43 ± 1.62 | 3.57 ± 0.15 |
|  | 20:4n-6 | 14(15)-EpETRE | 7.61 ± 1.26 | 1.71 ± 0.41 | 6.68 ± 1.17 | 5.14 ± 1.57 | 12.99 ± 2.84 | 4.56 ± 1.13 | 7.43 ± 1.06 | 6.50 ± 0.65 |
|  | 20:4n-6 | 5,6-DiHETRE | 0.03 ± 0.02 | 0.04 ± 0.01 | 0.14 ± 0.04 | 0.17 ± 0.07 | 0.08 ± 0.02 | 0.09 ± 0.03 | 0.14 ± 0.04 | 0.16 ± 0.04 |
|  | 20:4n-6 | 8,9-DiHETRE | 0.22 ± 0.02 | 0.13 ± 0.03 | 0.19 ± 0.02 | 0.20 ± 0.04 | 0.29 ± 0.05 | 0.18 ± 0.04 | 0.24 ± 0.05 | 0.15 ± 0.06 |
|  | 20:4n-6 | 11,12-DiHETRE | 1.09 ± 0.09 | 0.73 ± 0.14 | 0.80 ± 0.09 | 0.99 ± 0.05 | 1.46 ± 0.18 | 1.23 ± 0.12 | 1.37 ± 0.14 | 1.21 ± 0.17 |
|  | 20:4n-6 | 14,15-DiHETRE | 1.43 ± 0.12 | 1.15 ± 0.23 | 1.32 ± 0.14 | 1.25 ± 0.11 | 2.14 ± 0.28 | 1.88 ± 0.10 | 2.18 ± 0.20 | 2.07 ± 0.18 |
|  | 20:4n-6 | 20-HETE | 10.64 ± 1.55 | 3.40 ± 1.01 | 21.68 ± 4.16 | 20.62 ± 8.15 | 41.19 ± 9.54 | 13.02 ± 3.89 | 21.17 ± 3.84 | 1.90 ± 1.26 |
|  | 20:5n-3 | 8(9)-EpETE | 0.34 ± 0.04 | 0.11 ± 0.06 | 0.34 ± 0.06 | 0.25 ± 0.08 | 1.13 ± 0.31 | 0.35 ± 0.14 | 0.58 ± 0.17 | 0.05 ± 0.03 |
|  | 20:5n-3 | 11(12)-EpETE | 0.42 ± 0.12 | 0.04 ± 0.03 | 0.34 ± 0.13 | 0.21 ± 0.10 | 1.11 ± 0.25 | 0.14 ± 0.13 | 0.57 ± 0.29 | 0.14 ± 0.09 |
|  | 20:5n-3 | 14(15)-EpETE | 0.94 ± 0.41 | 0.37 ± 0.18 | 0.78 ± 0.16 | 0.97 ± 0.37 | 2.39 ± 0.45 | 1.21 ± 0.38 | 1.55 ± 0.46 | 0.89 ± 0.24 |
|  | 20:5n-3 | 17(18)-EpETE | 0.94 ± 0.13 | 0.29 ± 0.12 | 0.86 ± 0.11 | 0.65 ± 0.16 | 2.50 ± 0.41 | 1.04 ± 0.33 | 2.12 ± 0.22 | 0.46 ± 0.15 |
|  | 20:5n-3 | 18-HEPE | 1.61 ± 0.15 | 0.97 ± 0.24 | 2.16 ± 0.30 | 1.97 ± 0.36 | 7.68 ± 1.09 | 4.01 ± 0.62 | 5.91 ± 0.61 | 2.00 ± 0.57 |
|  | 22:6n-3 | 7(8)-EpDPE | 2.99 ± 0.18 | 1.44 ± 0.42 | 6.65 ± 1.80 | 4.29 ± 1.53 | 9.62 ± 2.20 | 2.87 ± 0.72 | 3.15 ± 0.43 | 0.73 ± 0.09 |
|  | 22:6n-3 | 10(11)-EpDPE | 11.12 ± 0.76 | 5.08 ± 1.50 | 27.05 ± 7.32 | 17.19 ± 6.49 | 39.07 ± 9.61 | 10.34 ± 2.27 | 12.45 ± 1.87 | 2.57 ± 0.24 |
|  | 22:6n-3 | 13(14)-EpDPE | 5.70 ± 0.45 | 2.78 ± 0.77 | 13.03 ± 3.32 | 30.49 ± 9.99 | 15.85 ± 3.90 | 5.66 ± 1.08 | 6.64 ± 0.93 | 1.89 ± 0.22 |
|  | 22:6n-3 | 16(17)-EpDPE | 4.19 ± 0.33 | 2.06 ± 0.53 | 9.25 ± 2.28 | 5.67 ± 1.85 | 14.21 ± 2.83 | 4.30 ± 0.72 | 5.03 ± 0.54 | 1.36 ± 0.13 |
|  | 22:6n-3 | 19(20)-EpDPE | 5.42 ± 0.29 | 2.90 ± 0.74 | 16.07 ± 4.76 | 8.47 ± 2.80 | 21.84 ± 3.39 | 12.11 ± 2.18 | 10.76 ± 0.54 | 4.76 ± 0.50 |
|  | Sum |  | 258.46 ± 15.40 | 78.03 ± 18.94 | 242.80 ± 39.38 | 200.17 ± 75.48 | 333.14 ± 53.84 | 158.50 ± 19.64 | 237.55 ± 27.35 | 98.98 ± 8.10 |

Values are mean ± SEM of 5-7 mice/group. ND = Below limits of detection of the assay.

**Supplemental Table 1D:** Specialized pro-resolving mediators (SPM) (pg/mg) in young and aged mouse tibialis anterior (TA) muscles

| Pathway | Substrate | Analyte | Uninjured |  | Day 1 post-injury |  | Day 3 post-injury |  | Day 5 post-injury |  |
| --- | --- | --- | --- | --- | --- | --- | --- | --- | --- | --- |
|  |  |  | Young | Aged | Young | Aged | Young | Aged | Young | Aged |
| Specialized pro-resolving mediators | 20:4n-6 | LXA4 | 0.16 ± 0.03 | 0.12 ± 0.04 | 0.22 ± 0.07 | 0.16 ± 0.04 | 0.55 ± 0.12 | 0.30 ± 0.09 | 0.35 ± 0.07 | 0.25 ± 0.06 |
|  | 20:4n-6 | LXA5 | ND ± ND | ND ± ND | ND ± ND | ND ± ND | ND ± ND | ND ± ND | ND ± ND | ND ± ND |
|  | 20:4n-6 | LXB4 | ND ± ND | ND ± ND | ND ± ND | ND ± ND | ND ± ND | ND ± ND | ND ± ND | ND ± ND |
|  | 20:4n-6 | 15-epi LXA4 | ND ± ND | ND ± ND | ND ± ND | ND ± ND | ND ± ND | ND ± ND | ND ± ND | ND ± ND |
|  | 20:4n-6 | 15-oxo LXA4 | ND ± ND | ND ± ND | ND ± ND | ND ± ND | ND ± ND | ND ± ND | ND ± ND | ND ± ND |
|  | 20:5n-3 | RvE1 | ND ± ND | ND ± ND | ND ± ND | ND ± ND | ND ± ND | ND ± ND | ND ± ND | ND ± ND |
|  | 20:5n-3 | RvE2 | ND ± ND | ND ± ND | ND ± ND | ND ± ND | ND ± ND | ND ± ND | ND ± ND | ND ± ND |
|  | 20:5n-3 | RvE3 | 0.39 ± 0.06 | ND ± ND | ND ± ND | ND ± ND | ND ± ND | ND ± ND | ND ± ND | ND ± ND |
|  | 22:5n-3 | RvD5(n-3DPA) | ND ± ND | ND ± ND | ND ± ND | ND ± ND | ND ± ND | ND ± ND | ND ± ND | ND ± ND |
|  | 22:5n-3 | PD1(n-3, DPA) | ND ± ND | ND ± ND | ND ± ND | ND ± ND | ND ± ND | ND ± ND | ND ± ND | ND ± ND |
|  | 22:5n-3 | MaR1(n-3DPA) | 0.64 ± 0.15 | 0.62 ± 0.09 | 1.41 ± 0.61 | 0.76 ± 0.14 | 1.66 ± 0.28 | 1.27 ± 0.19 | 1.64 ± 0.17 | ND ± ND |
|  | 22:6n-3 | RvD1 & AT-RvD1 | ND ± ND | ND ± ND | ND ± ND | ND ± ND | ND ± ND | ND ± ND | ND ± ND | ND ± ND |
|  | 22:6n-3 | 8-oxoRvD1 | 0.31 ± 0.05 | ND ± ND | 0.51 ± 0.07 | ND ± ND | ND ± ND | ND ± ND | ND ± ND | ND ± ND |
|  | 22:6n-3 | 17-oxoRvD1 | ND ± ND | ND ± ND | ND ± ND | ND ± ND | ND ± ND | ND ± ND | ND ± ND | ND ± ND |
|  | 22:6n-3 | RvD2 | ND ± ND | ND ± ND | ND ± ND | ND ± ND | ND ± ND | ND ± ND | ND ± ND | ND ± ND |
|  | 22:6n-3 | RvD3 | ND ± ND | ND ± ND | ND ± ND | ND ± ND | ND ± ND | ND ± ND | ND ± ND | ND ± ND |
|  | 22:6n-3 | AT-RvD3 | ND ± ND | ND ± ND | ND ± ND | ND ± ND | ND ± ND | ND ± ND | ND ± ND | ND ± ND |
|  | 22:6n-3 | RvD4 | ND ± ND | ND ± ND | ND ± ND | ND ± ND | ND ± ND | ND ± ND | ND ± ND | ND ± ND |
|  | 22:6n-3 | RvD5 | ND ± ND | ND ± ND | ND ± ND | ND ± ND | ND ± ND | ND ± ND | ND ± ND | ND ± ND |
|  | 22:6n-3 | RvD6 | 0.38 ± 0.03 | 0.37 ± 0.05 | 1.20 ± 0.30 | 1.03 ± 0.23 | 2.91 ± 0.44 | 1.29 ± 0.22 | 0.99 ± 0.12 | 0.70 ± 0.13 |
|  | 22:6n-3 | PD1 | ND ± ND | ND ± ND | 10.96 ± ND | 9.84 ± ND | 12.50 ± 4.17 | ND ± ND | 5.45 ± 0.54 | ND ± ND |
|  | 22:6n-3 | 10S,17S-DiHDoHE | ND ± ND | ND ± ND | ND ± ND | ND ± ND | 22.51 ± 6.32 | ND ± ND | 10.63 ± 1.15 | ND ± ND |
|  | 22:6n-3 | AT-PD1 | ND ± ND | ND ± ND | ND ± ND | ND ± ND | ND ± ND | ND ± ND | ND ± ND | ND ± ND |
|  | 22:6n-3 | 22-OH-PD1 | ND ± ND | ND ± ND | ND ± ND | ND ± ND | ND ± ND | ND ± ND | ND ± ND | ND ± ND |
|  | 22:6n-3 | Maresin1 | 4.04 ± 1.44 | 1.02 ± 0.49 | 10.38 ± ND | 1.48 ± ND | 9.58 ± 5.50 | 2.81 ± 1.07 | 3.92 ± 0.53 | ND ± ND |
|  | 22:6n-3 | 7(S)-Maresin1 | ND ± ND | ND ± ND | ND ± ND | ND ± ND | ND ± ND | ND ± ND | ND ± ND | ND ± ND |
|  |  | Sum | 6.26 ± 1.90 | 1.98 ± 0.54 | 5.41 ± 2.14 | 3.59 ± 1.77 | 32.19 ± 12.39 | 3.26 ± 0.60 | 14.12 ± 2.33 | 0.87 ± 0.19 |

Values are mean ± SEM of 5-7 mice/group. ND = Below limits of detection of the assay.

**Supplemental Table 1E:** Non-enzymatic metabolite concentration (pg/mg) in young and aged mouse tibialis anterior (TA) muscles

| Pathway | Substrate | Analyte | Uninjured |  | Day 1 post-injury |  | Day 3 post-injury |  | Day 5 post-injury |  |
| --- | --- | --- | --- | --- | --- | --- | --- | --- | --- | --- |
|  |  |  | Young | Aged | Young | Aged | Young | Aged | Young | Aged |
| Non-enzymatic | 20:4n-6 | 9-HETE | ND ± ND | ND ± ND | ND ± ND | ND ± ND | ND ± ND | ND ± ND | ND ± ND | ND ± ND |
|  | 20:4n-6 | 11-HETE | 25.77 ± 1.33 | 15.63 ± 3.44 | 31.22 ± 4.21 | 35.00 ± 7.79 | 118.40 ± 14.45 | 78.22 ± 18.00 | 111.29 ± 7.04 | 67.45 ± 8.69 |
|  | 20:5n-3 | 9-HEPE | 10.28 ± 2.03 | 5.13 ± 1.56 | 4.98 ± 0.94 | 3.93 ± 0.60 | 23.36 ± 3.67 | 16.56 ± 2.59 | 25.88 ± 3.59 | 12.56 ± 2.11 |
|  | 20:5n-3 | 11-HEPE | 1.46 ± 0.10 | 0.92 ± 0.24 | 1.81 ± 0.27 | 1.91 ± 0.41 | 8.64 ± 1.17 | 4.61 ± 0.94 | 8.08 ± 0.24 | 2.55 ± 0.61 |
|  | 22:6n-3 | 8-HDoHE | 5.59 ± 0.41 | 3.34 ± 0.81 | 20.91 ± 5.19 | 15.96 ± 6.61 | 35.77 ± 6.40 | 11.82 ± 2.56 | 13.73 ± 1.59 | 5.22 ± 0.52 |
|  | 22:6n-3 | 10-HDoHE | 8.03 ± 1.10 | 4.96 ± 1.19 | 20.09 ± 4.25 | 16.46 ± 6.11 | 36.87 ± 4.73 | 15.31 ± 2.42 | 17.33 ± 1.24 | 8.07 ± 0.91 |
|  | 22:6n-3 | 11-HDoHE | 3.71 ± 1.25 | 2.61 ± 0.50 | 14.22 ± 3.61 | 11.29 ± 4.56 | 24.36 ± 4.43 | 10.17 ± 1.86 | 10.70 ± 1.25 | 4.99 ± 0.51 |
|  | 22:6n-3 | 13-HDoHE | 6.30 ± 0.27 | 4.86 ± 0.99 | 20.30 ± 4.41 | 16.43 ± 5.68 | 41.85 ± 5.32 | 18.58 ± 3.50 | 22.17 ± 1.89 | 10.47 ± 1.16 |
|  | 22:6n-3 | 16-HDoHE | 8.89 ± 0.66 | 6.70 ± 1.38 | 28.65 ± 6.72 | 24.67 ± 9.70 | 54.86 ± 8.30 | 21.27 ± 3.39 | 24.13 ± 3.41 | 10.57 ± 1.29 |
|  | 22:6n-3 | 20-HDoHE | 8.09 ± 0.34 | 6.29 ± 1.20 | 24.33 ± 5.49 | 20.52 ± 6.13 | 53.26 ± 7.44 | 23.13 ± 3.61 | 22.98 ± 1.78 | 12.11 ± 1.25 |
| Sum |  |  | 78.12 ± 3.14 | 50.45 ± 10.31 | 166.52 ± 33.30 | 146.18 ± 46.42 | 397.38 ± 50.58 | 199.69 ± 37.72 | 256.31 ± 13.38 | 134.00 ± 14.98 |

Values are mean ± SEM of 5-7 mice/group. ND = Below limits of detection of the assay.

